# Transcriptomic remodeling of the retina in a Zebrafish model of Retinitis Pigmentosa

**DOI:** 10.1101/2022.10.04.510882

**Authors:** Abirami Santhanam, Eyad Shihabeddin, Haichao Wei, Jiaqian Wu, John O’Brien

**Affiliations:** Department of Ophthalmology & Visual Science, McGovern Medical School, The University of Texas Health Science Center at Houston; University of Houston College of Optometry; MD Anderson UT Health Graduate School of Biomedical Sciences; Department of Neurosurgery, McGovern Medical School, The University of Texas Health Science Center at Houston

**Author notes:** Address for correspondence: Abirami Santhanam, John O’Brien, Department of Vision Science, University of Houston College of Optometry, 4401 Martin Luther King Blvd, JDA 2195, Houston, TX 77204. Contributed equally (Abirami Santhanam, Eyad Shihabeddin).

## Abstract

Inherited retinal degenerative diseases such as Retinitis Pigmentosa (RP) result in progressive loss of photoreceptors until an individual is completely blind. A hallmark of these diseases is progressive structural and functional remodeling of the remaining retinal neurons as rod photoreceptors are lost. While many studies focus on regenerative or bionic therapies to restore vision, extensive remodeling of retinal cell types throughout the course of retinal degenerative diseases stands as a barrier for successful implementation of these strategies. As a window onto the molecular basis of remodeling, we have performed a comparative analysis of single-cell transcriptome data from adult Zebrafish retina of wild-type and a P23H mutant rhodopsin model of RP. In addition to providing a benchmark atlas of retinal cell type transcriptomes in the wild-type adult Zebrafish retina, we find transcriptional changes in essentially all retinal cell types in the P23H model. Increased oxidative stress is evident not only in the rods but also in cones, retinal pigmented epithelium (RPE) and to a lesser extent in amacrine and bipolar cells. Metabolic changes increasing oxidative metabolism and glycolysis are found in rods and cones, while evidence of increased activity of the mitochondrial electron transport chain is found in retinal ganglion cells (RGCs). Evidence of synaptic remodeling is found throughout the retina, with changes to increase synaptic transmission in photoreceptors and bipolar cells, increased ionotropic glutamate receptors in amacrine and ganglion cells, and dendritic and axon remodeling throughout. Surprisingly, RPE, cones and bipolar cells in the P23H retinas also have increased expression of genes involved in circadian rhythm regulation. While this model system undergoes continuous regeneration, ongoing remodeling impacts the entire retina. This comprehensive transcriptomic analysis provides a molecular road map to understand how the retina remodels in the context of chronic retinal degeneration with ongoing regeneration.

## Introduction

Retinitis Pigmentosa (RP) is a debilitating, genetically-based disease that affects 1 in 4000 people worldwide [1–3]. Early symptoms of RP include loss of peripheral and night vision due to the death of rod photoreceptors [4], generally progressing to complete blindness due to the death of cone photoreceptors via a bystander effect [5, 6]. Symptoms can begin in young adults with progressive loss of vision; some patients are completely blind by their early thirties [7]. Gene therapy for one variant of RP has been somewhat effective [8], but due to the large number of genes that can cause RP [9], there are no treatment options for most patients. Regenerative medicine and stem cell transplantation offer hope to combat the progression to blindness and preserve vision [10, 11].

A potential barrier to implementing regenerative and transplantation strategies is the extensive remodeling of retinal cell types that are not themselves directly affected by the mutations that cause degeneration [12–15]. Along with remodeling comes aberrant spontaneous neuronal activity, obscuring the transmission of visual signals [16–18]. Understanding the molecular underpinnings of this remodeling will be paramount to developing successful strategies to regenerate functional retina.

The revolution in large-scale genomic and transcriptomic methods has revealed the transcriptional diversity of cell types with unprecedented detail. These techniques have been used to resolve cell type complexity in the retina [19–21] and to examine development [22]. Recent studies have revealed the retinal response to acute injury, including the transcriptional changes that support the regeneration of neurons from Müller glial precursors [23]. However, these studies lack the context of retinal remodeling found in chronic retinal degenerative diseases. We have previously characterized a transgenic Zebrafish line expressing P23H mutant rhodopsin, which models chronic Retinitis Pigmentosa with ongoing photoreceptor degeneration and regeneration [24]. In the present study, we have performed single-cell RNA Sequencing (sc-RNA Seq) of adult retina to assess the transcriptomic remodeling that takes place in the P23H Zebrafish model. In addition to providing a benchmark atlas of retinal cell type transcriptomes in the wild-type adult Zebrafish retina, we find transcriptional changes in essentially all cell types, reflective of the remodeling that has gone on in the chronic disease. Prominent changes reveal pathology in retinal pigmented epithelium (RPE) structural integrity, synaptic remodeling in several cell types, and disruption in circadian clock gene expression. We have further identified the retinal progenitor cell population differentiating into rod photoreceptors. Collectively, our results provide a comprehensive understanding of the transcriptomic changes that take place throughout the retina in a model with chronic retinal degeneration. These findings will provide the field of vision restoration with a more precise atlas of genes unique and specific to each cell type in the retina. Ultimately, this will allow for therapeutic approaches to become more cell-and gene-specific.

## Results

### Single-cell transcriptomic atlas of Zebrafish retina

To generate a cell atlas of the adult Zebrafish retina, we obtained six retinas from three wild-type AB strain fish (WT) at the age of 7 months. The retinas were collected along with RPE and single cells were obtained as described in detail in the materials and methods section. We analyzed them with single◻cell RNA sequencing (scRNA◻seq) using the 10X Genomics Chromium platform. We have repeated the experiments twice with different single-cell chemistries V2 and V3. Both runs provided similar cluster patterns and cell types. The data presented here are from the V3 chemistry as this has improved Median Genes per Cell and Median UMI Counts (unique transcripts) per Cell. Following quality control and filtering using the Seurat package [25], as described in detail in the methods section, the final dataset contained 13,551 cells, which were taken forward for further analysis. The scRNA◻seq data were initially analyzed using an unsupervised graph clustering approach implemented in Seurat (V3) to classify individual cells into cell populations according to similarities in their transcriptome profiles. Finally, we used the tSNE plot to visualize clusters in a two-dimensional space. Overall, the cells were classified into 23 transcriptionally distinct clusters (Fig. 1A).

**Figure 1.**
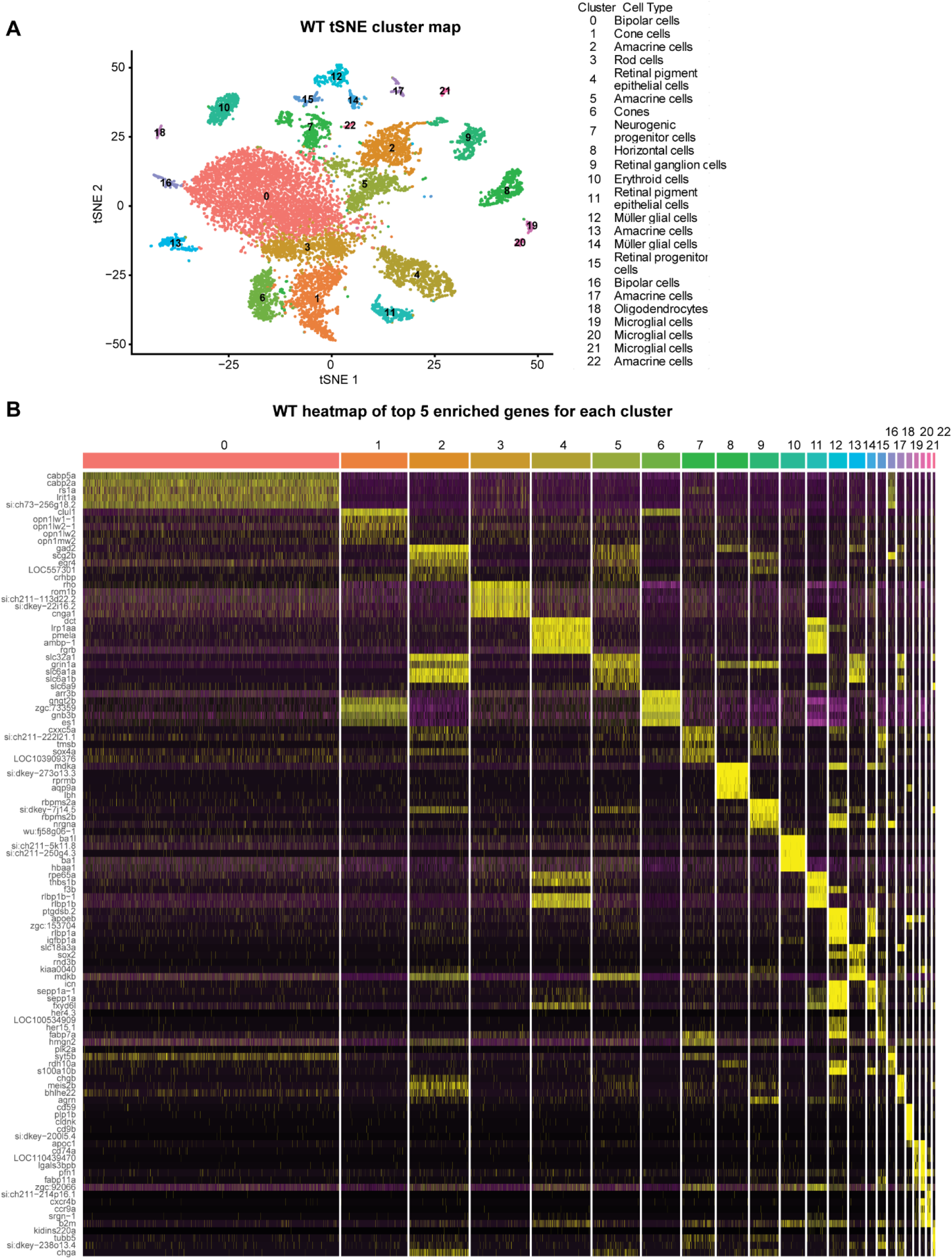
Single-cell RNAseq analysis on Zebrafish retina. A) tSNE dimensional reduction and visualization of transcriptional profiles of 13,551 retinal cells from WT zebrafish. B) Heatmap showing the top 5 genes enriched for each cluster.

The clusters were identified into different retinal cell types using markers identified in previous studies (Table S1). We were not able to assign any cellular identity to one cluster as it expressed markers from multiple retinal cell types. This cluster is not shown in the figure and was not used for further analysis. We have identified all the major retinal cell types including five clusters of amacrine cells, three clusters of microglia, two clusters each of cones, bipolar cells, retinal pigmented epithelium (RPE) and Müller glia, and one cluster each of rods, horizontal cells, retinal ganglion cells, neurogenic progenitor cells (NPCs), oligodendrocytes and retinal progenitor cells (RPCs). Each cell type cluster expresses a unique set of genes along with some shared genes between clusters that define it as a specific cell type. The top 5 genes that are specifically enriched in each cell type are displayed in the heatmap shown in Fig. 1B; a list of the top 10 genes in each cluster is listed in Table S2.

### Cellular changes in a Zebrafish model of Retinitis Pigmentosa

To identify transcriptional changes that occur during chronic retinal degeneration and regeneration, we performed the same type of analysis on a single cell transcriptome library prepared from the adult retina of a transgenic Zebrafish model of Retinitis Pigmentosa expressing P23H mutant rhodopsin [24]. Following quality control and filtering using the Seurat package [25] as described in the methods section, our final dataset contained 15,511 cells.

We compared the WT and P23H datasets by using an integration algorithm, which aims to identify shared cell states that are present across different datasets [26] (Fig. 2A). Using the integrated analysis, we detected all the major cell types that are present in the WT dataset, but there were differences in the number of cells in different cell types (Fig.2A, 2B). There was a reduction in the number of mature rod photoreceptors (cluster 3) in the P23H retina, reflecting the ongoing degeneration occurring in this model system. In addition, a class of newly differentiated rods (cluster 13) that shows both progenitor and rod markers that did not appear in the clustering of the WT dataset was present. While some cells from the WT retina fell into this cluster, this class was 3-fold more abundant in the P23H retina, reflecting the continuous regeneration underway in the P23H retina (Fig. 2B, 2C). In keeping with this, we also detected a 4-fold higher number of retinal progenitor cells (RPCs) in the P23H retina compared to the WT (Fig.2C, cluster 14). Interestingly we saw an 8-fold decrease in retinal pigmented epithelial cells (RPE, clusters 10 and 15) compared to WT (Fig.2C).

**Figure 2.**
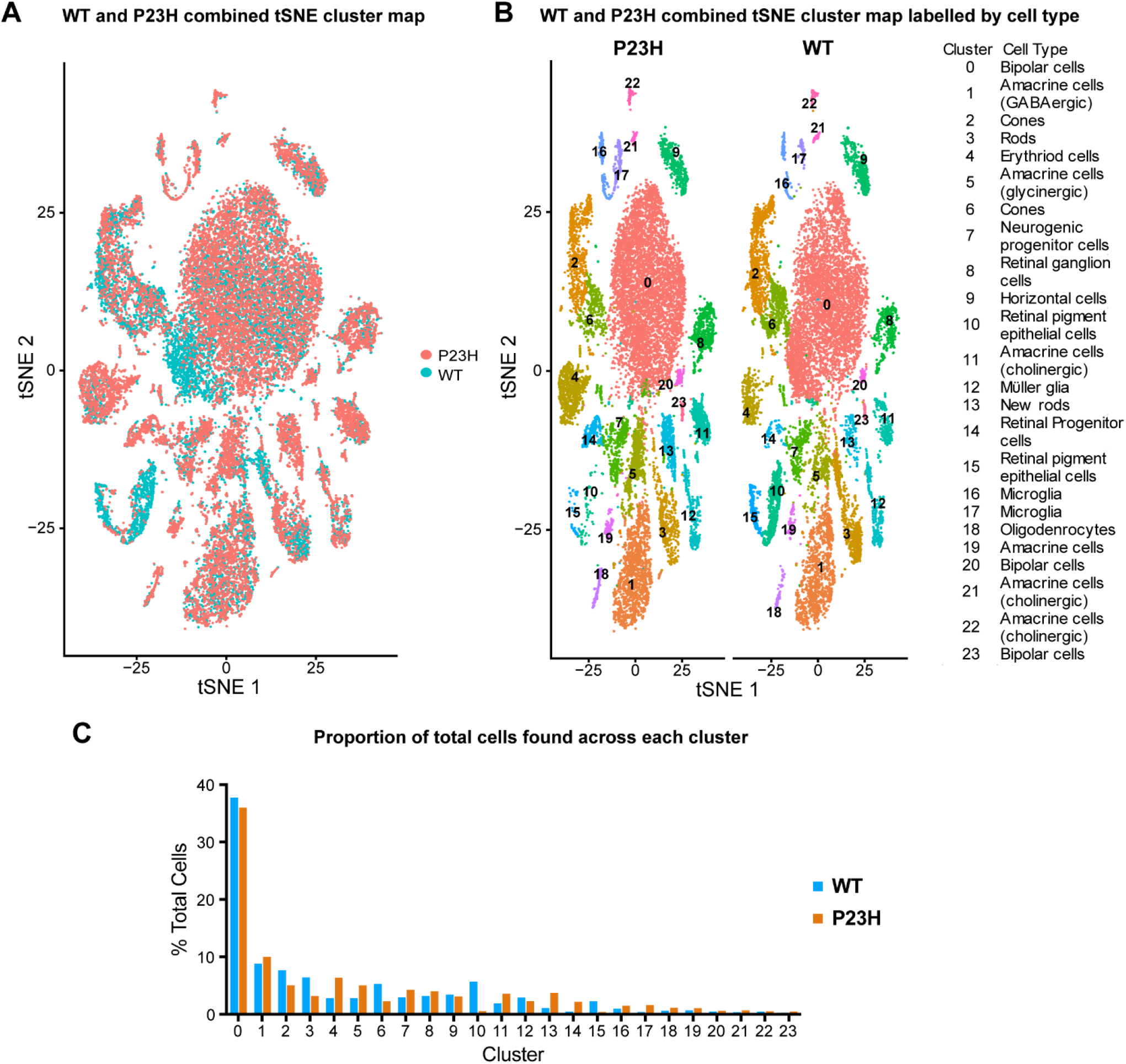
Single-cell RNAseq analysis on WT-P23H integrated Zebrafish retina dataset. A) Overlap of tSNE dimensional reduction and visualization of single-cell transcriptome from WT and P23H transgenic fish B) tSNE showing the proportions of all the cell types found in the P23H and WT retinal samples. C) Bar graph showing the proportions of all the cell types found in the P23H and WT retinal samples.

Numerical comparisons of cell numbers in single-cell transcriptome data present challenges for interpretation. Cells of different types are not captured with equal efficacy and tissue dissociation conditions can favor the preservation of some cell types and loss of others (e.g. the larger number of bipolar cells than photoreceptors in our datasets). To reinforce the estimates of relative changes in numbers of cells of certain types, we also examined the numbers of cells in an earlier dataset using 10x V2 chemistry prepared using different animals of comparable age. Table S3 shows that trends in the relative ratios of cell numbers are conserved between datasets. In this dataset, we find that rods are more than 2-fold reduced in the P23H whereas the newly-formed rods are 9-fold higher in P23H. Our dataset from V2 chemistry also showed a reduction of RPE cells in P23H compared to the WT (2.7 fold decrease in P23H) (Table S3).

### Rod photoreceptors display enhanced oxidative metabolism, oxidative stress and synaptic remodeling

Our previous study showed that rod photoreceptors undergo continuous cell death and that the number of rods is almost 4-fold less in the P23H retina than in the WT [24]. Our current single-cell study also reflects the decrease in the number of rods in the P23H dataset compared to the WT (Fig. 3A). Violin plots of expression of individual genes in a cluster allow us to compare gene expression levels in P23H and WT conditions. Figure 3B shows violin plots of the rod dominant genes rhodopsin (*rho*) and rod arrestins (*saga* and *sagb*). The expression of all of these genes was conserved between WT and P23H. We performed functional pathway analysis using Cystoscope Cluego [27] and Metascape [28] analysis software (Fig. S1). The P23H rod dataset revealed an increased level of markers for oxidative stress (*sirt2, atf4a, oxr1b*), response to misfolded proteins (*hsp70.3, hspa5, dnajb1b, xbp1*), actin depolymerization (*eps8, sptbn1*) (Fig. 3C), and lipid phosphorylation (*dgkzb, pik3ca, pik3r3a*) (Fig. 3D), reflecting physiologic responses to the P23H mutant rhodopsin that ultimately lead to photoreceptor degeneration. Interestingly we have also seen the increased transcriptome of certain ciliary proteins (*cep290, bbs5, rpgrip1, rp1*) and calcium channels (*cacna1c*, *tmco1*) (Fig. 3D).

**Figure 3.**
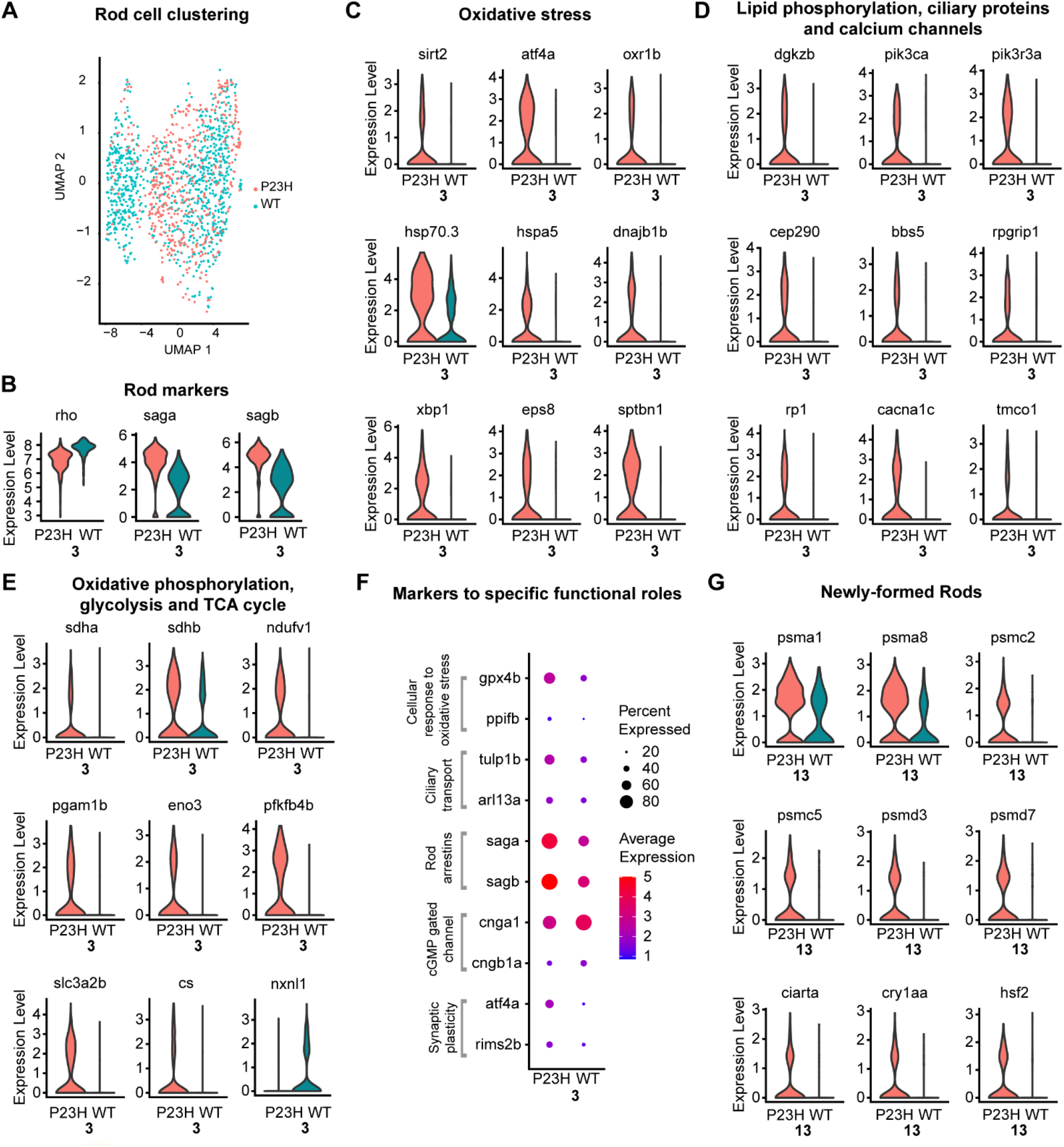
Transcriptomic analysis of rod cells and newly formed rod cells in the WT-P23H integrated dataset reveal increased response to oxidative stress and misfolded proteins. A) Overlap of tSNE dimensional reduction and visualization of rod cell transcriptome from WT and P23H transgenic fish B) Violin plots showing levels of expression of canonical marker genes in WT and P23H rod cells. C), D), and E) Violin plots showing levels of expression of DEGs involved in different functional pathways including oxidative stress, response to misfolded proteins, metabolism, synaptic remodeling, etc., between WT and P23H rod cells. F) Dot plot showing the expression level and the percentage of cells of DEGs involved in different functional pathways between WT and P23H. G) Violin plots showing levels of expression of DEGs involved in proteasome and ubiquitin-mediated degradation pathway and stress response markers between WT and P23H rod cells.

Metabolism-related differentially expressed genes (DEGs) were upregulated in the P23H rods including oxidative phosphorylation (*sdha*, *sdhb*, *ndufv1*), glycolysis (*pgam1b*, *eno3*, *pfkfb4b*, *slc3a2b*) and TCA cycle (*cs* and *ndufv1*) (Fig. 3E). It is possible that the stress due to misfolded protein response leads to increased energy demand that is met by increased oxidative metabolism. Notably, *nxnl1*, the Zebrafish homolog of Rod-derived Cone Viability factor, is strongly decreased in P23H rods (Fig. 3E). This factor promotes glucose uptake in cones [29] and potentially rods [30]; its loss could result in reduced glucose uptake. Our pathway analysis depicts the increase in pathways involved in the synthesis of precursor metabolites, glycolysis, pyruvate metabolism and TCA cycle to cope with increased stress and energy demand in the P23H rods (Fig. S1). It has been shown recently by a multi-omic study that early metabolic imbalance and mitochondrial stress in neonatal photoreceptors lead to cell death in the Pde6b rd1 mouse model of retinal degeneration [31]. Our results correlate with the metabolic changes reported in that study. Importantly we noticed an increase in *gpx4b* and *ppifb*, which are involved in the cellular response to oxidative damage, suggesting that rods are experiencing oxidative stress. DEGs involved in ciliary transport including *tulp1b* and *arl13a* are also increased in P23H. Mutations in *tulp1* are reported to contribute to ~5% of total RP cases [32, 33] and mutations in *arl13* lead to failure of rod outer segment formation [34, 35]. Interestingly we noted a significant increase in the transcript level of rod arrestins (*saga*, *sagb*) in P23H compared to the WT (Fig. 3B, F). The arrestins may be upregulated to compensate for the constitutive activation of P23H mutant rhodopsin as it has shown previously that in the mammalian retina stable complexes are formed between mutant rhodopsin and rod arrestin leading to rod cell death [36]. It is implicated that tulp1b is responsible for the ciliary transport of arrestins [37] and the increased arrestins may accumulate in cilia leading to cell death. DEGs involved in cyclic nucleotide-gated channels including *cnga1* and *cngb1a* are both reduced in P23H. Both are involved in the phototransduction cascade and their decrease suggests a defect in phototransduction [38]. The DEGs involved in the regulation of synaptic plasticity (*atf4a*, *rims2b*) are upregulated in the P23H, suggesting changes in the synaptic region of P23H rods (Fig. 3F). Previous studies in Zebrafish models of rod damage do not reveal such comprehensive transcriptional changes [39–42] and it is important to understand these changes at the molecular level to target for right therapeutic interventions.

### A newly-forming rod cell cluster is evident in P23H

We identified cluster 13 as newly-forming rods, as this cluster expresses both progenitor cell markers and rod photoreceptor markers. The progenitor markers including *stmn1a*, *ccnd1* and *hes*6 are present at a low level, whereas the rod differentiation markers including *fabp7a*, *cxxc5a*, *arl13a* and *neurod1* are present at high levels in the transcriptome, representing the shift of cells from progenitor state to a differentiated state. This cluster of newly-forming rods was not evident in the WT dataset when clustered alone (Fig. 1A), and can be only seen in WT after integrating with P23H for the cluster analysis (Fig. 2A, B). In keeping with the persistent loss and regeneration of rods in the P23H animals, there was a larger number of newly-forming rods in P23H (3.8% of cells in the dataset) compared to the WT (1.1% of cells). Functional analysis with ClueGo and Metascape showed that pathways involved in the proteasome and ubiquitin-mediated degradation (UPR), eukaryotic translation, rod cell development, and eye morphogenesis are enhanced in the P23H transcriptome (Fig. S1C). The transcripts of genes involved in proteasome and ubiquitin-mediated degradation including *psma1*, *psma8*, *psmc2*, *psmc5*, *psmd3*, *psmd7*, and stress response markers including *ciarta*, *cry1aa* and *hsf2* are more highly expressed in the P23H dataset compared to WT (Fig. 3G), suggesting that even newly forming rods are experiencing stress associated with expression of the P23H mutant rhodopsin.

### Cone photoreceptors show markers of stress and changes in metabolic and circadian genes

We initially identified two cone clusters, cluster 2 and cluster 6. The short wavelength opsin *opn1sw1* is seen in cluster 6 and the long-wavelength opsin *opn1lw1* is enriched in cluster 2 (Fig. 4A). While expression of the P23H mutant rhodopsin is limited to the rods [24], cones in the P23H retina show an increase in the transcriptome of *hsf2* associated stress response pathway (Fig. 4B), suggesting that cones in this model experience elevated physiological stress. There may be increased oxidative stress in cones due to increased exposure to oxygen in the absence of fully functioning rods, which may lead to increased *hsf2* and subsequent activation of the MAPK signaling pathway [43]. Our data further confirmed the increased expression of *mapk6* in P23H cones compared to WT (Fig. 4B). Cones in the P23H retina also show increased expression of genes for glycolysis and the TCA cycle (*pfkfb3*, *pfkfb4b*, *eno3*, *cs*, *aco2*, *got1*) (Fig. 4B), but like rods, expression of *nxnl1* is reduced in cones. These results imply that cones rely on aerobic metabolism, but that glucose supply could be limiting. It has been shown previously that aerobic glycolysis in photoreceptors is essential for normal rod function and is a metabolic choice that augments cone function and survival during nutrient stress conditions [44]. We also noticed increased expression in DEGs involved in ribose phosphate biosynthesis (*socs3s*, *gcdha*) and RNA polymerase II transcription and elongation (*cdk7*, *polr2a*, *gatad2ab*) (Fig. 4B). Sufficient supplies of NADPH, ATP and the metabolic intermediates ensure rapid macromolecular synthesis underlying the continuous self-renewal of cone OS and the demand may be increased to cope with the increased stress.

**Figure 4.**
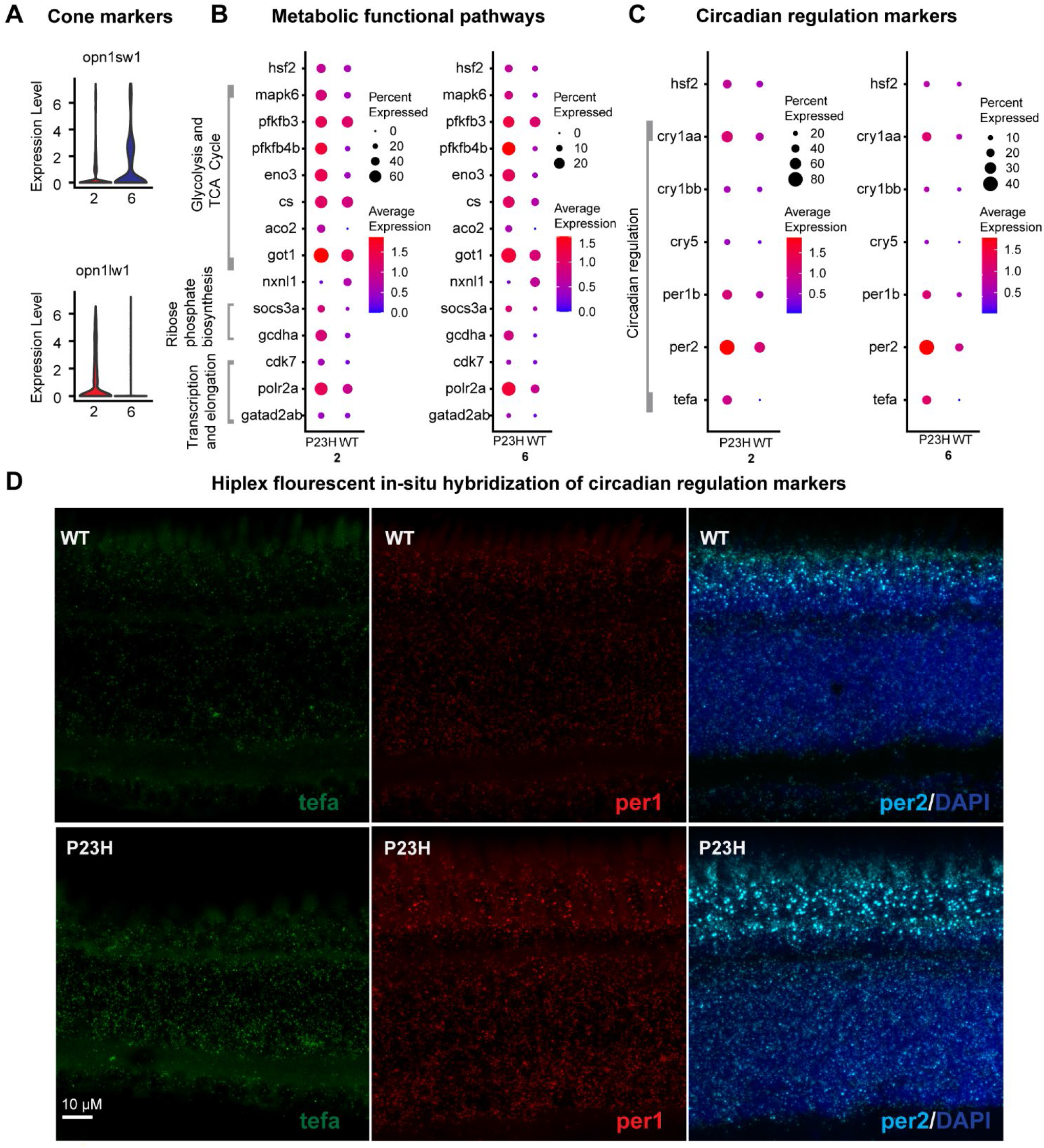
Comparative transcriptomic analysis of cone cells between WT-P23H integrated dataset reveals changes in metabolism and circadian regulation. A) Violin plots showing levels of expression of short and long wavelength opsin leading to cone clusters 2 and 6 in the integrated transcriptome. B) Dot plot showing the expression level and the percentage of cells of DEGs involved in different metabolic functional pathways between WT and P23H. C) Dot plot showing the expression level and the percentage of cells of DEGs involved in circadian regulation between WT and P23H. D) Hiplex in-situ fluorescent hybridization images for tefa, per1, and per2 show increased expression in P23H compared to WT reflecting the changes captured by single-cell transcriptome analysis.

The *hsf2*-associated stress response pathway includes increased levels of *cry1a* and *nr1d2b*, which are shown to be involved in circadian function [45]. It has been shown that in Zebrafish larvae the expression of *hsf2* was concomitant with the expression of genes involved in the response to oxidative stress and chaperone genes, and it occurs in phase with *cry1a* and other genes that belong to the negative arm of the transcriptional regulation network of the circadian rhythm [46, 47]. We examined the transcriptome profile of other molecular circadian regulators and found that the circadian regulators *cry1aa*, *cry1bb, per1b*, *per2* and *tefa* were increased in the P23H cones compared to the WT (Fig. 4C). To validate the single-cell transcriptome data, we examined the expression of *per1*, *per2* and *tefa* using multiplex in situ hybridization. The results demonstrate the increased expression *per1*, *per2* and *tefa* in the outer nuclear layer (ONL) of the P23H retina compared to the WT (Fig. 4D). Disruption of behavioral and retinal circadian rhythms has recently been reported in a mouse model of RP due to an RNA splicing factor mutation [48], suggesting commonality in pathology. Overall these changes suggest that the cones undergo comprehensive modifications including metabolism and circadian activity during the degeneration/regeneration scenario.

### RPE tight junction genes and stress-protective mechanisms are lowered in the RP model

The Zebrafish retinal transcriptome showed two different RPE clusters, clusters 10 and 15 (Fig. 5A). Both RPE cell clusters express the canonical RPE genes *rpe65a*, *dct* and *lrp1aa* (Fig. 5B) but are segregated due to the presence of transcriptome that enriches for different functional pathways (Fig. S2). Differential expression analysis revealed that cluster 15 has many unique transcripts that were not expressed in cluster 10 (Fig. 5C). The 5 most up-regulated genes in RPE cluster 10 were RNA binding proteins including *rpl3, rpl4, rpl5a* and *rpl6*, which enable protein translation; these were also present in cluster 15. Functional pathways involved in phagocytosis, endocytosis, lysosome, oxidative stress and drug metabolic process are all enriched in cluster 15, whereas cluster 10 showed pathways involved in translation and ribosome assembly (Fig. S2). The receptor tyrosine kinase *mertka* involved in RPE phagocytosis of photoreceptor outer segments (POS) [49, 50] is highly expressed in cluster 15, suggesting active phagocytosis in these RPE cells. It has been shown that the structure and function of the RPE vary depending on the location in the retina [51, 52], consistent with the distinct populations we detect in the single-cell transcriptome data.

**Figure 5.**
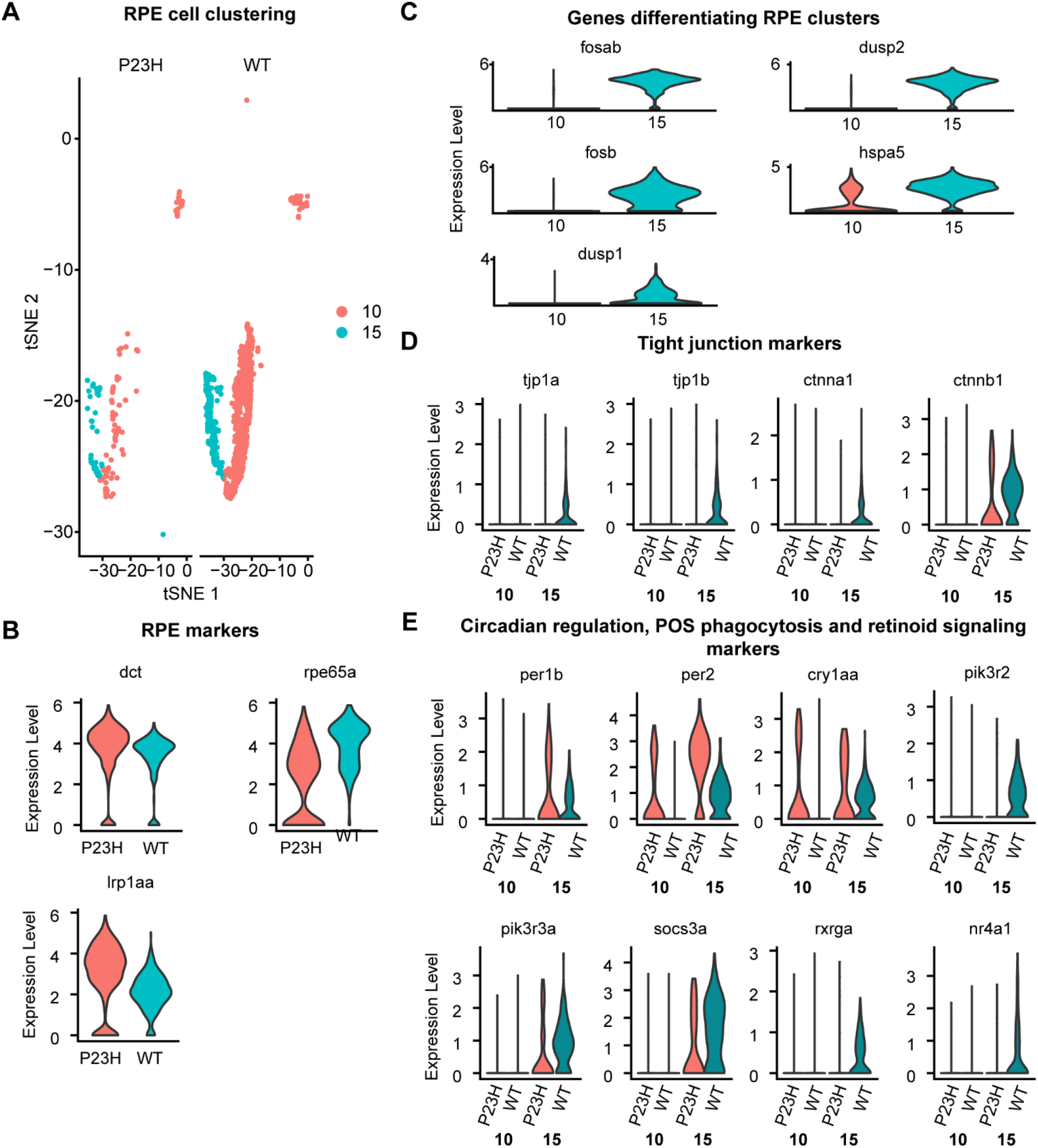
RPE tight junction transcriptome is decreased in the RP model. A) tSNE showing the proportions of RPE cell CLUSTERS found in the P23H and WT retinal samples B) Violin plots showing levels of expression of canonical marker genes in WT and P23H RPE cells. C) Violin plots showing levels of expression of DEGs between two RPE clusters 10 and 15. D) Violin plots showing levels of expression genes involved RPE tight junction between WT and P23H cells. E) Violin plots showing levels of expression of DEGs including those involved in circadian regulation, POS phagocytosis, and retinoid signaling between WT and P23H RPE cells.

We have seen a drastic decrease in the number of RPE cells in the P23H RP model compared to the WT (Fig 5A), which is consistent with several previously reported studies that when the rods are lost oxygen continues to flow into the outer retina from the choriocapillaris, creating a hyperoxic environment that is presumably hostile for the remaining cells [53, 54]. In some retinal degeneration models, disruption of RPE tight junctions is observed, affecting the integrity of RPE cells [15, 55]. Analysis of the DEGs in RPE cells involved in tight junction formation, including *tjp1a*, *tjp1b*, *ctnna1* and *ctnnb1*, revealed a decrease in the P23H compared to the WT (Fig. 5D). This suggests that RPE tight junction disruption is a pathological effect in RP that is conserved across species.

Ongoing degeneration and regeneration in the P23H RP model had a significant impact on several other aspects of the RPE transcriptome. We have noticed an increase in circadian genes such as *per2*, *per1b* and *cry1aa* in the P23H dataset (Fig 5E), similar to that seen in cones, suggesting the change is coordinated between RPE and photoreceptors. It has been shown in previous studies that *per2* regulates POS phagocytosis in RPE cells [47], suggesting a change in the rhythm of POS phagocytosis in the RP model. The phosphoinositide 3-kinase regulators *pi3kr2, pi3kr3aA* and *socs3a* are reduced in the P23H retina compared to the wild-type (Fig. 5E). It has been shown in previous studies that the activation of the PI3K-Akt pathway protects RPE cells against the deleterious effects of oxidative stress that occur due to POS phagocytosis [56, 57]. These protective mechanisms seem to be reduced in the RPE of P23H Zebrafish.

The retinoid X receptor *rxrga* and nuclear receptors *nr4a1* and *nr4a3* are lowered in the P23H dataset compared to the WT (Fig. 5E). It has been shown that NR4A receptors heterodimerize with RXR receptors and activate transcription in a 9-cis retinoic acid-dependent manner. They can also repress inflammatory gene promoters by recruiting corepressor complexes [58, 59]. All these transcriptome changes suggest that the RPE cells undergo high dynamic changes during the degeneration-regeneration scenario and lose several stress protective mechanisms.

### Rod bipolar cells show evidence of stress and neuronal remodeling

We have identified three different bipolar cell clusters, 0, 20 and 23 in our dataset; each cluster expressed a specific set of markers (Fig. 6A). The rod bipolar cell marker *prkca* [19, 26] is highly enriched in cluster 20 (avg. expression 2.7) and to a minor extent in cluster 0 (avg. expression 0.7) only in very few cells (Fig. 6B). Because of the high expression of *prkca*, we considered cluster 20 likely to be the rod-dominated mixed bipolar cell [60] that forms most synaptic contacts with rods. All of the bipolar cell clusters displayed differences in transcriptome profile between the WT and P23H (Fig. 6B), including downregulation of neuronal specification genes *neurod4* and *nrxn3b* and upregulation of stress-related genes including *ubl3a*. A prominent change in all bipolar cell clusters is the dramatic upregulation of *rgs16* (Fig. 6B), which accelerates the offset of G-protein signaling [61, 62], potentially modifying signaling by metabotropic glutamate receptors, dopamine receptors or other GPCRs.

**Figure 6.**
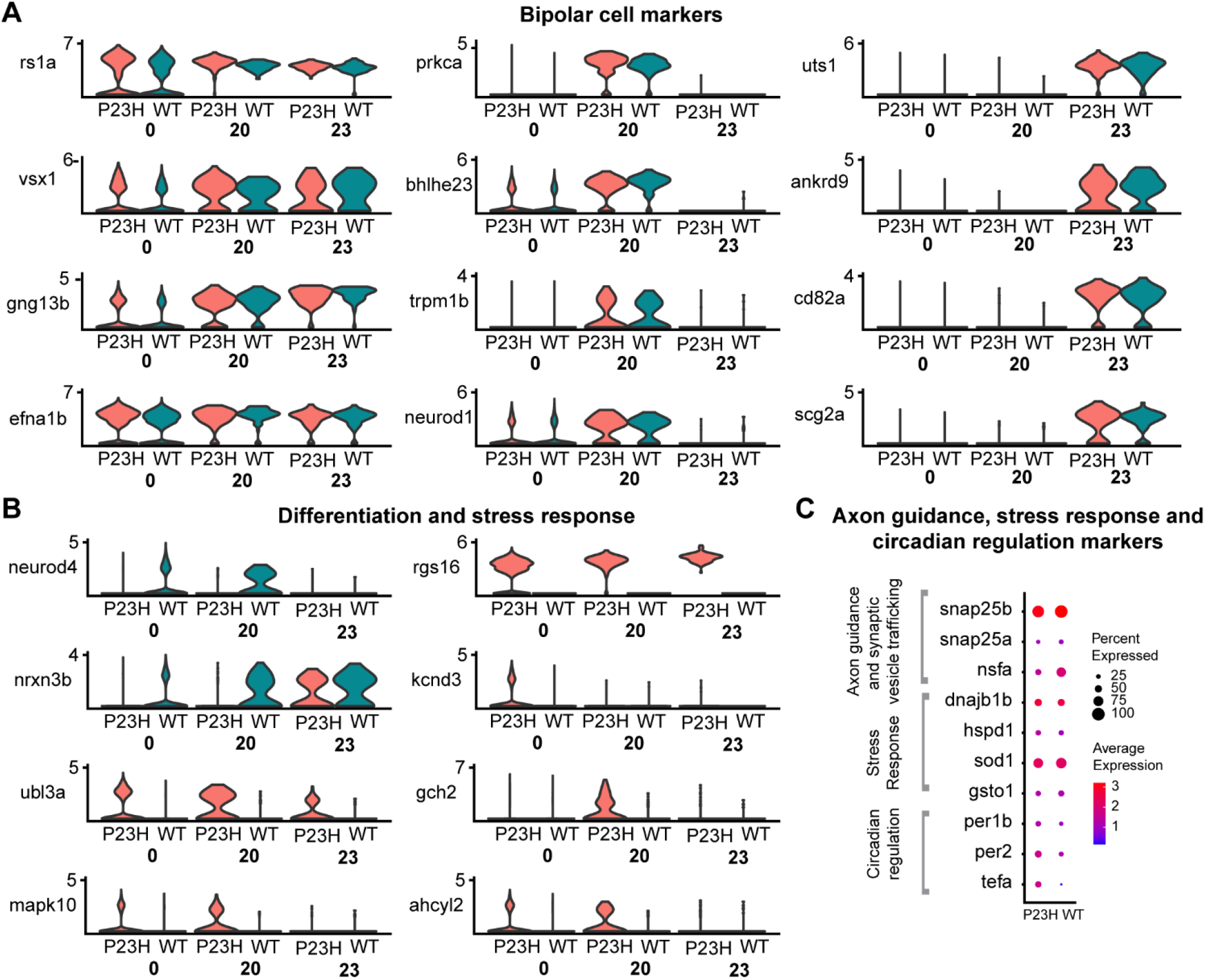
Comparative transcriptomic analysis of bipolar cell clusters in the WT-P23H integrated dataset shows evidence of stress and neuronal remodeling in the RP model. A) Violin plots showing levels of expression of specific marker genes in different clusters of bipolar cells (0, 20, and 23). B) Violin plots showing levels of expression of DEGs between the three bipolar cell clusters. C) Dot plot showing the expression level and the percentage of cells of DEGs involved in axon guidance, stress response, and circadian regulation between WT and P23H.

We focused on cluster 20 to examine the influence of rod degeneration and regeneration on gene expression. Genes involved in synaptic vesicle trafficking including *snap25a, snap25b* and *nsfa* are increased in the P23H dataset compared to the WT (Fig. 6C). This suggests that there may be compensation at the bipolar cell output synapses for the reduced rod input in P23H retina. We also detected increases in the transcriptome of some genes involved in the stress response including *dnajb1b, hspd1, sod1* and *gsto1* in the P23H dataset compared to the WT (Fig. 6C). As we found in cones and RPE, levels of *per1b*, *per2* and *tefa* that are involved in circadian regulation were elevated (Fig. 6C). This suggests that rod photoreceptor loss leads to circadian changes that are reflected throughout the retina.

The overall comparison between WT RBP transcriptome and P23H RBP transcriptome in the STRING interaction network suggests that P23H RBP cells have much less interaction network compared to WT RBP (Fig. S3A). The top enriched pathways are different between WT and P23H (Fig. S3B). Pathways for ATP synthesis and reactive oxygen species (ROS) and reactive nitrogen species (RNS) production were enriched in the P23H RBP transcriptome, suggesting increased energy demand and stress response in the P23H RBP cells (Fig. S3B). Thus, physiological remodeling in the P23H RBP cells involves a complex interplay of oxidative stress, synaptic remodeling and synaptic transmission.

### Horizontal and amacrine cells show changes in genes regulating axonal remodeling in the P23H RP model

We identified the horizontal cells using the markers *rprmb, aqp9a* and *mdka* (Fig. 7A). The gap junction proteins *cx52.6, cx52.9 and cx55.5* are all uniquely enriched in the horizontal cells (Fig. 7B). We studied the differences between the horizontal cells in WT and P23H by functional analysis (Fig. S4) and identified increased expression of genes involved in axon remodeling and microtubule stability including *apc2, camsap1b, sptbn1, sqstm1*, *ndel1a* and zgc*:92606* (Fig. 7A). Previous studies have shown that *apc2* and *camsap1* play an essential role in axonal projections through the regulation of microtubule stability [63, 64]. Structural remodeling of horizontal cells, including neurite sprouting, is a well-documented phenomenon in models of retinal degeneration [65, 66], in keeping with these findings. Functional annotation of genes differentially expressed in WT and P23H horizontal cells show response to nitrogen starvation GO enriched in P23H suggesting an increase in autophagy and organelle disassembly (Fig. S4), indicating ongoing cellular stress.

**Figure 7.**
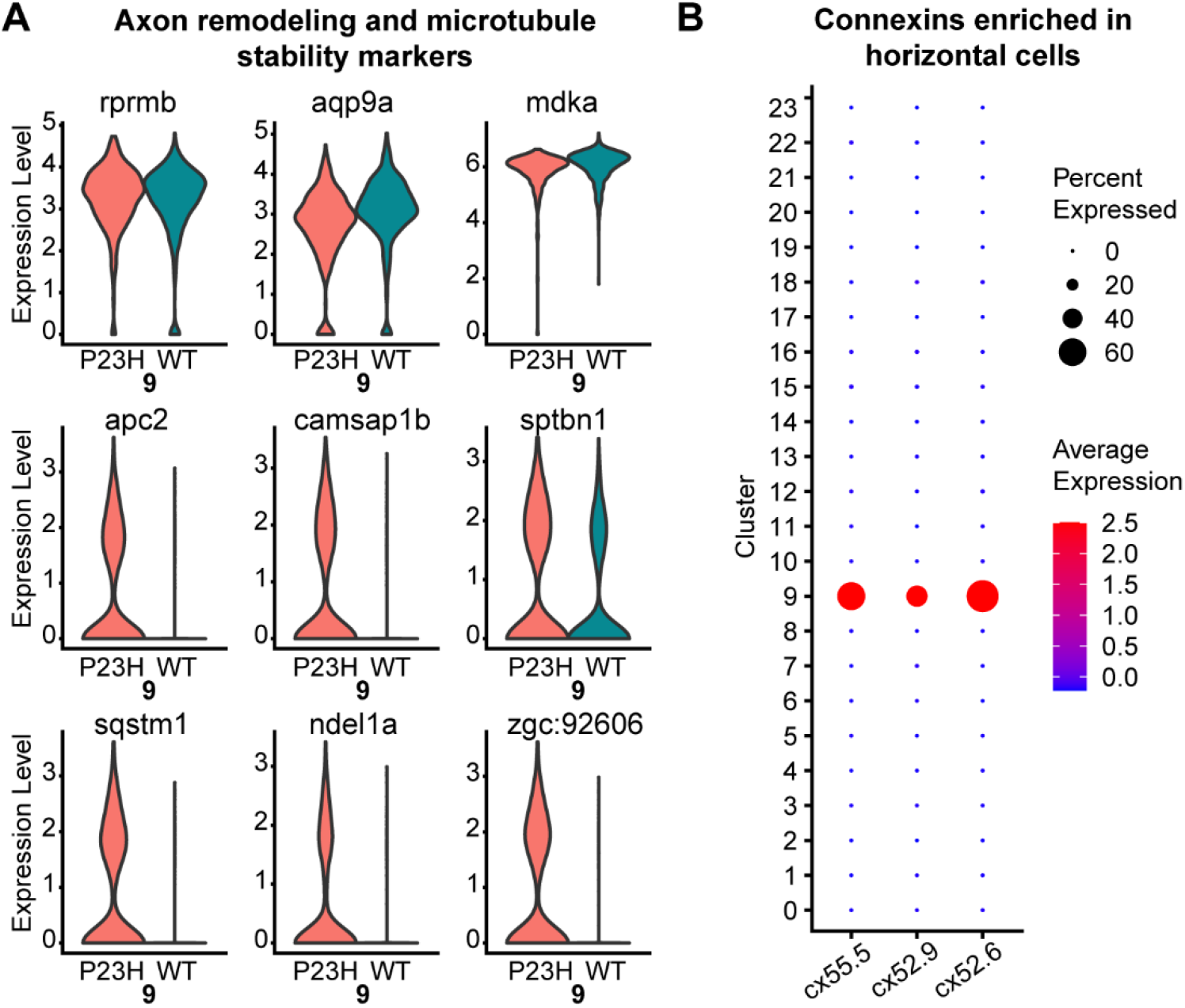
Comparative transcriptomic analysis of Horizontal cells in the WT-P23H integrated dataset reveals changes in axon remodeling in the RP model. A) Violin plots showing levels of expression of canonical marker genes as well as DEGs involved in axon remodeling and microtubule stability in WT and P23H horizontal cells. B) Dot plot showing the specific enrichment of *cx52.6*, *cx52.9*, and *cx55.5* in horizontal cell cluster (9) compared to all other retinal cell types.

We have identified six clusters of amacrine cell types with our initial clustering analysis including clusters 1, 5, 11, 19, 21 and 22 (Fig. 2B). Cluster 1 was identified as GABAergic amacrine cells due to the specific enrichment of the *gad2*, which is involved in GABA neurotransmitter biosynthesis (Fig. 8A). The glycinergic amacrine cells were identified by the enrichment of glycine transmembrane transporter *slc6a9* in cluster 5. Cluster 11 was identified as starburst amacrine cells due to the enrichment of both *gad2* and *chata*. Cluster 19 contained the unique enrichment of *calb1* and *calb2a* without the presence of glycine or GABA transporters (Fig. 8A). In the past studies have reported this kind of unique amacrine subpopulation [67] and it is interesting to see these data captured at a single-cell level even without enrichment of amacrine cells. This amacrine subset also shows the high expression of *nxph2a*, which is a signaling molecule that resembles neuropeptides [68]. Clusters 21 and 22 both express cholinergic markers *chata*, *slc18a3a* along with *chgb*, and are differentiated in that cluster 22 is GABAergic (expressing *gad2*) while cluster 21 is glycinergic (expressing *slc6a9*; Fig. 8A). Both of these clusters also share high similarities with the retinal ganglion cell (RGC) transcriptome. All six amacrine cell clusters are present in both WT and P23H.

**Figure 8.**
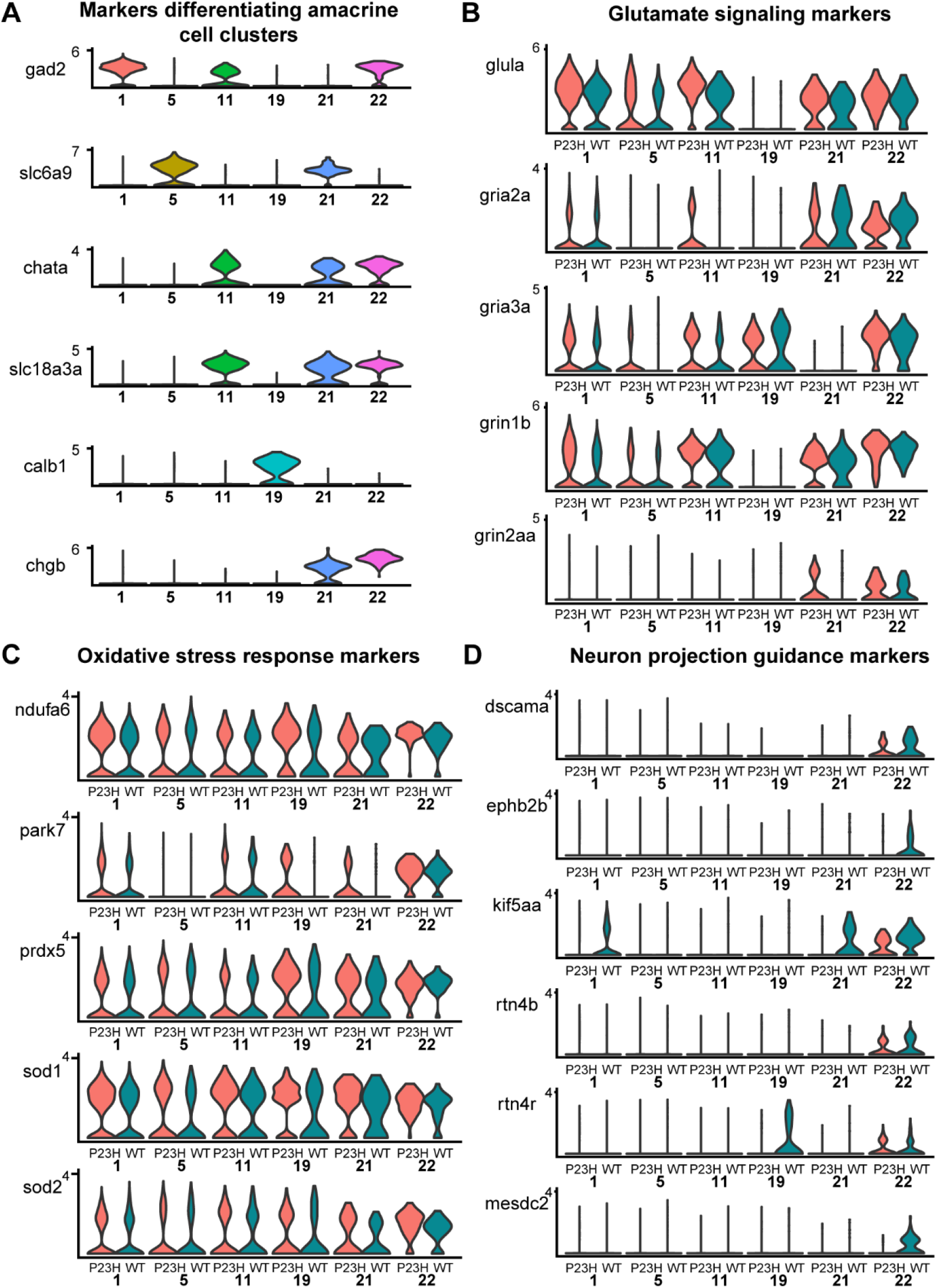
Transcriptomic analysis of Amacrine cells reveals six different amacrine cell clusters and changes in glutamate signaling and stress response among clusters. A) Our data identified six different types of amacrine clusters (1, 5, 11, 19, 21, and 22). Violin plots showing levels of expression of canonical marker genes expressed in different amacrine clusters. B), C), D) Violin plots showing levels of expression of DEGs involved in glutamate signaling (B), oxidative stress response (C), and neuron projection guidance (D) between WT and P23H amacrine cells respectively.

Changes in gene expression in amacrine cells were relatively modest between WT and P23H retina. We see the increased expression of *glula* in amacrine cell clusters in P23H. Further, the ionotropic glutamate receptors *gria2a, gria3a, grin1b and grin2aa* are all increased in P23H amacrine cell clusters compared to WT, suggesting the possibility of increased excitatory current (Fig 8B). We noticed small increases in genes that respond to oxidative stress and ROS, including *ndufa6, park7, prdx5, sod1* and *sod2* (Fig. 8C). We also detected a reduction in the expression of a few genes involved in neuron projection guidance [69] including *dscama, ephb2b, kif5aa, rtn4b*, *rtn4r* and *mesdc2* (Fig. 8D), suggesting defective neuronal wiring in P23H amacrine cells.

### Retinal ganglion cells show disturbances in the normal synaptic activity in P23H transgenic fish

Retinal ganglion cells in animal models of RP are characterized by aberrant spontaneous activity [16–18]. We identified the RGCs through the enriched presence of the markers *pou4f1, rbpms2a* and *rbpms2b*, identifying clusters 8 and 12 (Fig. 9A). In keeping with increased spontaneous activity observed in other RP models, pathway analysis of cluster 8 showed that certain genes involved in the regulation of synaptic transmission, especially glutamatergic synapses including *grin2aa, shank2* and *cacng2a*, are enriched in the P23H dataset compared to the WT (Fig. 9B). Further analysis revealed the increase of glutamate receptors including *grin1b*, *grin2aa*, *gria2a*, *gria2b* and *gria3a* as well in the P23H retina, suggesting changes in sensitivity to glutamatergic input due to photoreceptor degeneration (Fig. 9C). These markers are differentially expressed genes associated with glioma in previous studies [70]. Transcripts of genes involved in ATP hydrolysis including *atp1b1b* and atp*1a3a, atp5a1, atp5l* and *atp2a2b* are increased in the P23H dataset (Fig. 9B). We noticed increased expression of transcripts involved in succinate dehydrogenase complex and electron transport chain including *sdha*, *sdhb*, *sucla2* and *suclg1*, suggesting the remodeling due to photoreceptor degeneration leads to metabolic changes, possibly in response to increased energy demand (Fig. 9B). Heat shock protein *hspd1* (HSP60) is highly enriched in the P23H RGC cells compared to the WT (Fig.9D) and may reflect the role of this protein as a cellular defense mechanism in response to stress. The increases in expression of genes supporting responses to glutamatergic input and increased spontaneous activity observed in other RP models suggest that RGCs are at risk for glutamate excitotoxicity. It has been shown in a previous study that the HSP60 levels are high in the RGCs of glaucomatous eyes compared to normal eyes [71]. Finally, we also noticed an increase in Wnt signaling components in the P23H retina compared to the WT including *kras*, *foxo3a*, *rac3b* and *dctn2* (Fig.9D). It has been shown in previous studies that Wnt signaling regulates several aspects of RGC biology, including differentiation, proliferation, and axonal outgrowth [72].

**Figure 9.**
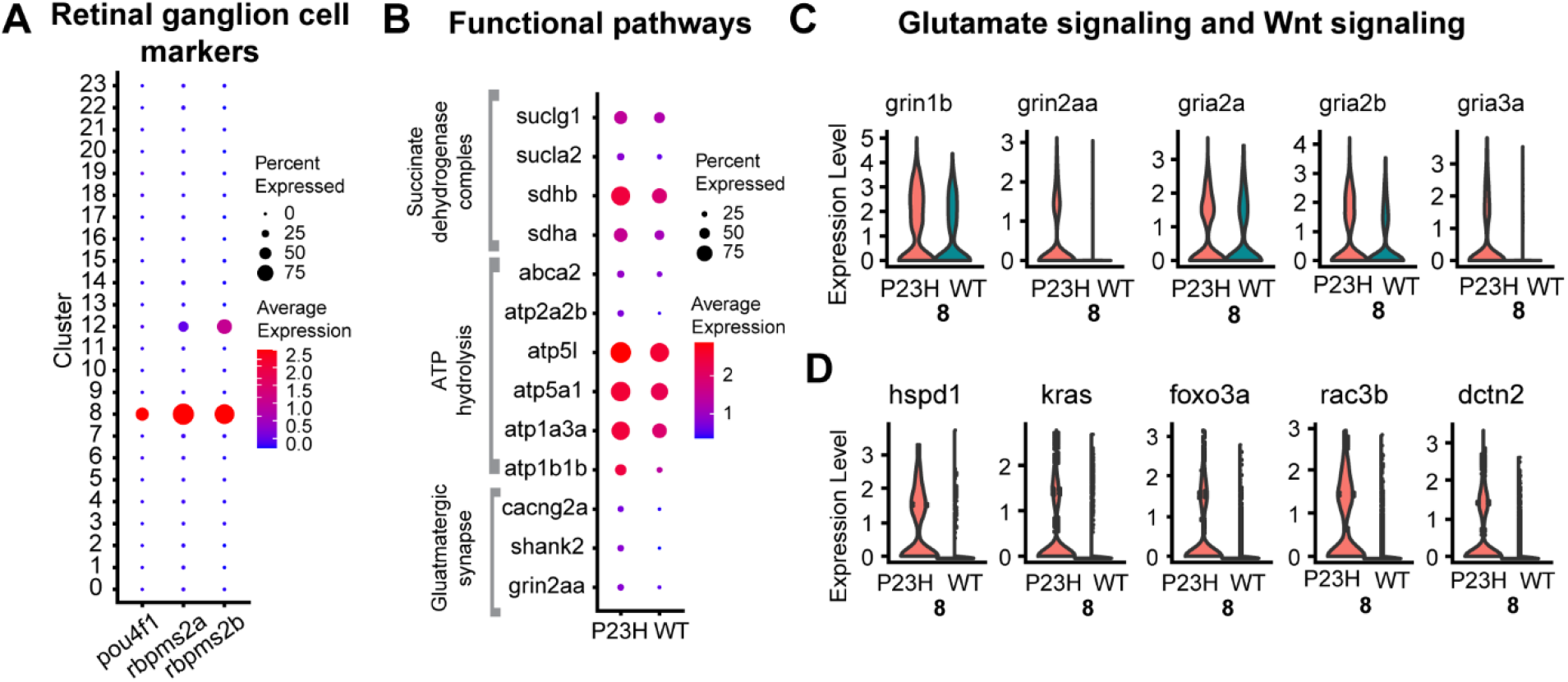
Comparative transcriptomic analysis of Retinal ganglion cells in WT-P23H integrated dataset. A) Dot plot showing the specific enrichment of canonical markers *pou4f1*, *rbpms2a*, and *rbpms2b* in the RGC cell cluster compared to all other retinal cell types. *rbpms2a* and *rbpms2b* are present at lower levels in Müller glial cells (12) as well. B) Dot plot showing the expression level and the percentage of cells of DEGs involved in different functional pathways between WT and P23H. C), and D) Violin plots showing increased levels of expression of DEGs involved in glutamate signaling (C) and Wnt signaling (D) between WT and P23H RGCs respectively.

### Phagocytic and apoptotic regulators are increased in P23H microglia/macrophages

We identified two microglial/macrophage clusters in the Zebrafish retina dataset, clusters 16 and 17, using the markers *cd74*a, *cd74bB* and *pfn1* (Fig. 10A). Cluster 16 showed the enriched presence of many markers including *apoc1*, *ccl34b.1* and *fabp11a*, whereas cluster 17 had the enriched presence of *ccl36.1*, *trac* and *zap70* (Fig. 10B). A recent study has reported the presence of two phenotypically and functionally distinct microglial populations in adult Zebrafish, segregated by the presence of *ccl34b.1* [73]. We differentiate the microglial clusters based on the presence of *apoc1* and *apoc1* (+)ve cluster 16 also shows the specific presence of *ccl34b.1* in our study. Another interesting study in the Zebrafish retina has identified the regeneration-associated transcriptional signature of retinal microglia and macrophages using RNA seq and most of the top 50 markers discussed in that study are captured in our single-cell analysis along with the specific type of microglia [74].

**Figure 10.**
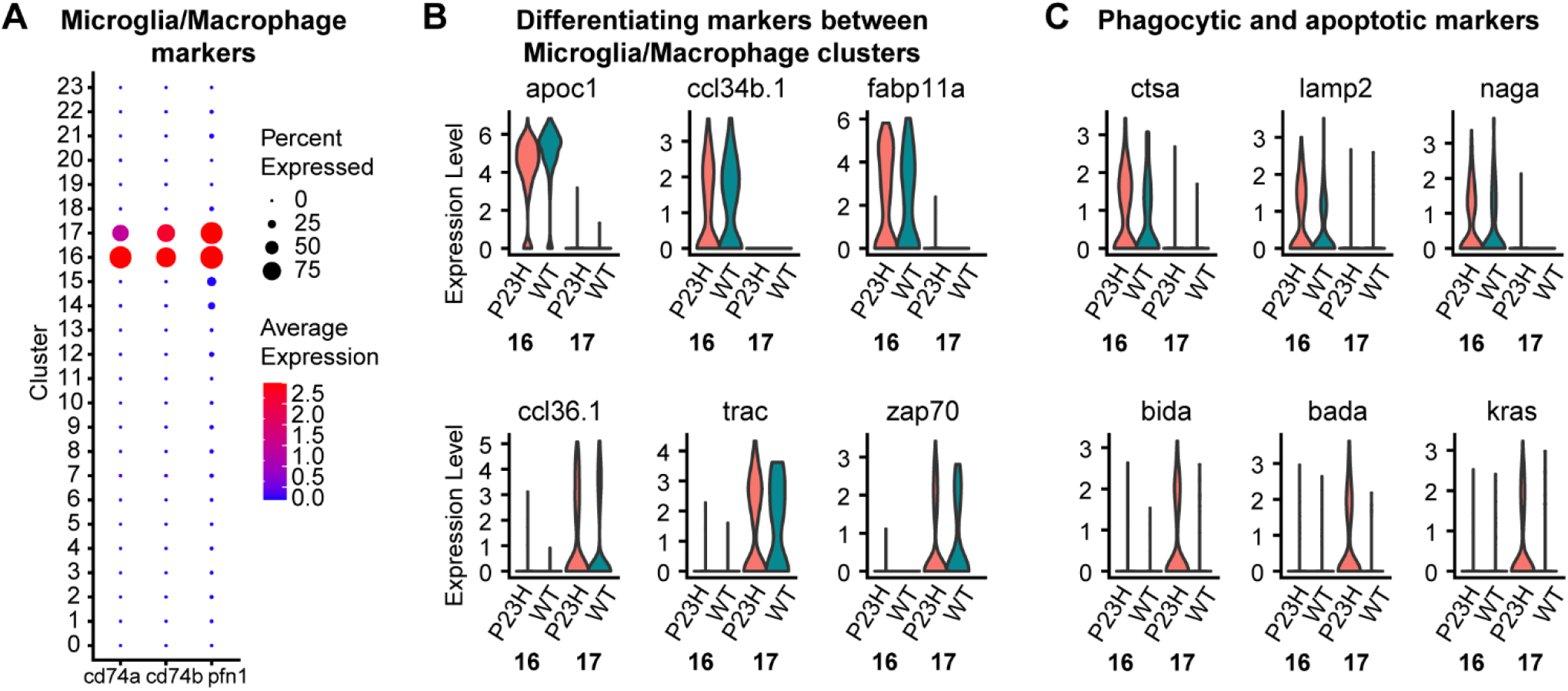
apoc1 (+) and apoc1 (−) microglial/macrophage clusters are identified in the Zebrafish retina. Dot plot showing the specific enrichment of canonical markers *cd74a*, and *cd74b* as well as the non-canonical marker *pfn1* in the microglia/macrophage cell clusters. B) Violin plots showing the expression of cluster-specific markers between the two microglial/macrophage clusters (16 and 17). (C) Violin plot showing the expression of increased phagocytic and apoptotic markers in P23H microglia/macrophages.

Functional analysis revealed differences between two microglial clusters: cluster 16 enriched with *apoc1* shows increased phagocytic and lytic activity along with cell chemotaxis and translation elongation, whereas cluster 17 shows the annotation for apoptosis, myeloid cell homeostasis, and regulation of the immune system (Fig S5). In the P23H fish certain phagocytic markers like *ctsa*, *lamp2* and *naga* are increased in cluster 16 whereas we see the increase of *bida*, *bada* and *kras* involved in the positive regulation of apoptosis in cluster 17 (Fig. 10C), suggesting functional changes in the microglial population during the degeneration-regeneration process in the retina. Our data provide several new molecular markers associated with microglia.

### Müller glia show increased proliferative function in the P23H RP model

Müller glia are the center of attention in studies examining the regeneration of neurons in the retina. Following retinal injury, Zebrafish Müller glia can reprogram to become multipotent progenitor cells with the ability to differentiate into many types of neuron [23, 75, 76]. We identified the Müller glia cluster 12 with the highly enriched expression of markers *ptgdsb.1*, *ptgdsb.2, aqp1a.1* and *rlbp1a* (Fig. 11A). Several transcriptional changes suggest that the Müller glial cells in the P23H chronic degeneration/regeneration model are tending towards their proliferative role and downregulating genes involved in the normal maintenance functions. Functional analysis showed enhanced TOR signaling and chromatin assembly or disassembly pathways in P23H whereas sterol biosynthesis and regulation of lamellipodium assembly were downregulated relative to the WT (Fig. S6). Further analysis of the transcriptome showed the increased expression of mTOR pathway regulators *sesn1* and *sik2b* in the P23H dataset (Fig. 11A); *synpo*, an actin-associated protein that modulates dendritic spine shape, is also enriched in the P23H Müller glial cells. We also noted an increase in *hmgcl* and *prodha* involved in the amino acid catabolic process in the P23H (Fig. 11A). Interestingly we also see a uniquely enriched expression of *hyal6* (hyaluronoglucosaminidase 6) in the P23H Müller glial transcriptome (Fig. 11A). It has been shown previously in Zebrafish studies that shorter forms of hyaluronic acid contribute to the regenerative process in both larval and adult Zebrafish [77, 78]. In the meantime, we see the unique reduction of markers for stem cell population maintenance including *notch1b*, *hmgcs1* and *snx5* in the P23H dataset compared to WT (Fig. 11B). We likewise see the decreased expression of genes involved in lamellipodium assembly such as *add3a, capzb* and *auts2b* as well as in sterol biosynthesis, including *adh5*, *cyp51* and *rdh10a* (Fig. 11B) in the P23H Müller glial transcriptome. Thus, Müller glia in the chronic retinal degeneration/regeneration model have shifted their transcriptional landscape to be poised to support proliferation.

**Figure 11.**
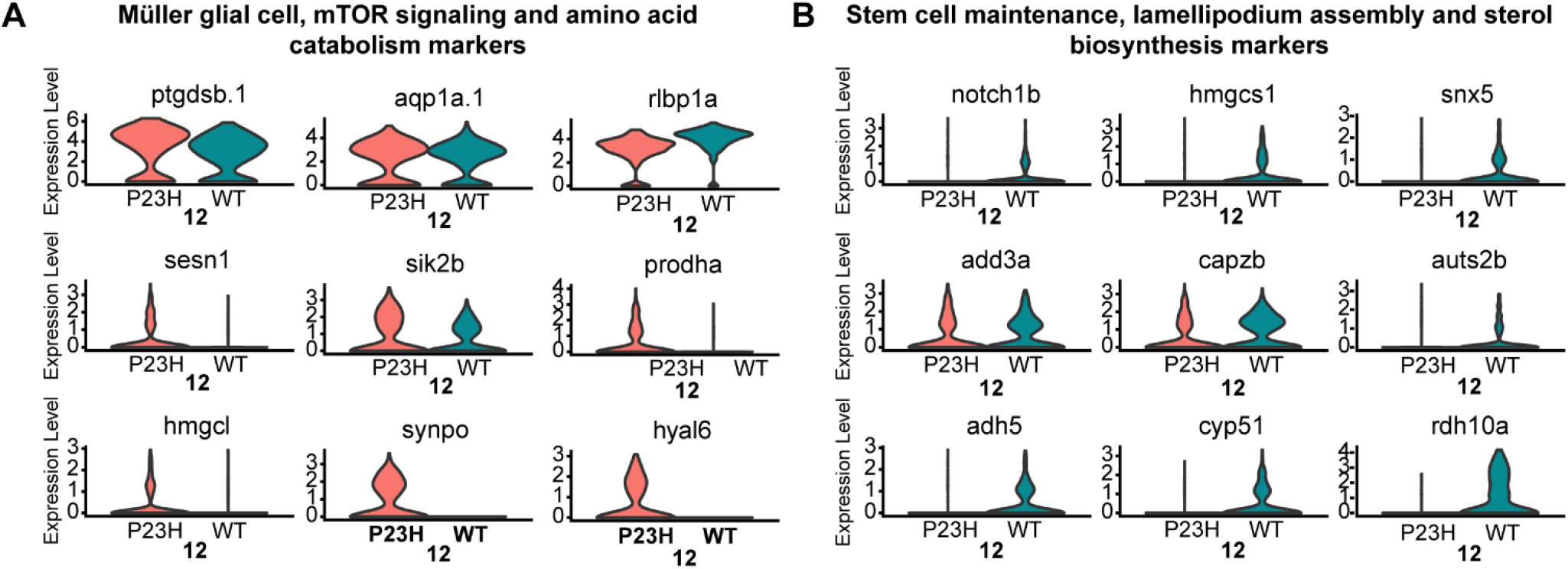
Müller glial cells in the P23H RP model show increased proliferative function. A) Violin plot showing the specific enrichment of canonical Müller glial cell markers as well as changes in DEGs involved in mTOR signaling as well as amino acid catabolism. B) Violin plot showing the decreased levels of DEGs involved in stem cell maintenance, lamellipodium assembly, and sterol biosynthesis.

### Retinal progenitor cells (RPCs) especially the rod progenitors are enriched in the P23H RP Zebrafish

The RPC population (cluster 14) was identified by the expression of proliferative stem cell markers including *ccna2, ccnb2, ccne2* and *stmn1a* (Fig. 12A). The P23H dataset contains a highly enriched amount of transcripts specific for mitotic cell division including *aurkb*, *top2a* and *ube2c*, suggesting a highly proliferative cell cluster in the P23H, whereas the WT transcriptome shows the enrichment of markers for pluripotent retinal progenitor cells including *foxn4*, *crabp2a*, *notch1b* and *her15.1* (Fig. 12A). It is very important to note that we identified a cluster of RPCs even in the wild type, suggesting that neurogenesis in Zebrafish is a very common phenomenon carried out to maintain the dynamic balance in the growth of eye size related to its body size. This also shows the power of single-cell profiling, which can identify small populations of cells in the tissue that might not be noticed in other types of analysis. The RPC population in WT is much smaller (~62 cells) compared to the P23H (~341 cells). Functional pathway analysis shows the enrichment of Notch signaling, FoxO signaling pathway, and TGF-B signaling pathway in the WT RPCs, whereas mitotic cell cycle and proteasome pathway, DNA replication, and chromosome organization are all enriched in P23H RPCs, suggesting a difference in the functional role of RPCs between WT and P23H retina (Fig. 12B).

**Figure 12.**
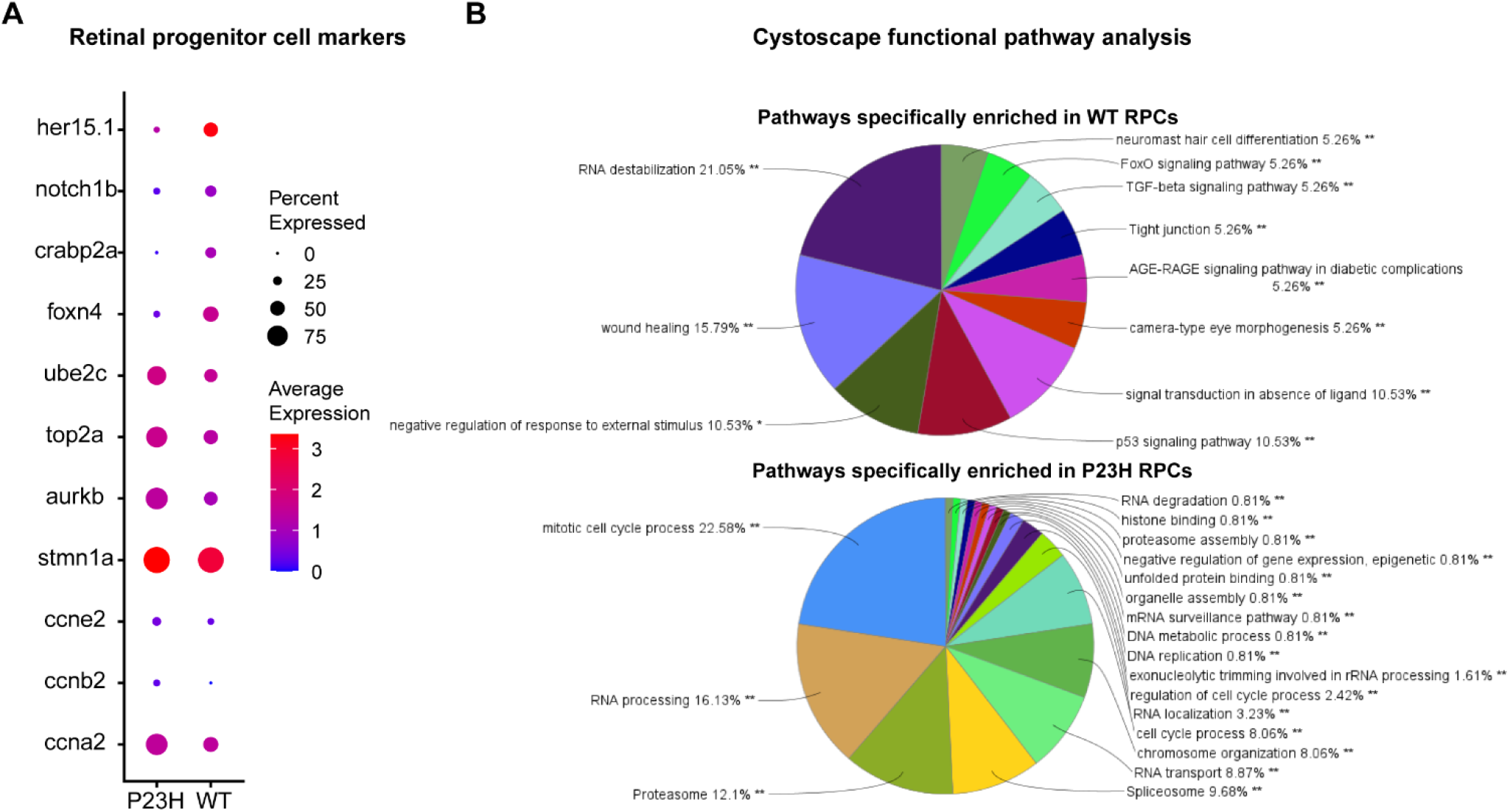
Retinal progenitor cells (RPCs) are enriched in the RP model. A) Dot plot showing the expression level and the percentage of WT (*her 15.1*, *notch1b*, *crabp2a*, *foxn4*) and P23H specific (*ube2c*, *top2a*, *aurkb*) RPC markers. B) Cytoscape functional pathway analysis generated pie charts showing the functional pathways that are specifically enriched in WT vs P23H RPCs.

### Neurogenic progenitor cells are identified in both WT and P23H Zebrafish retina

We identified a separate group of neurogenic progenitor cells (NPCs), cluster 7, which resemble committed neuronal cells but show the presence of several progenitor markers including *marcksl1a*, *marcksl1b*, *atoh7* and *myca*. It has been reported recently that two transcriptionally distinct progenitor clusters including the proliferative and neurogenic progenitors are seen in developing Zebrafish retina [23]. Our data confirms that this clusters are seen also in the Zebrafish adult retina supporting the continuous growth of Zebrafish eye. This cluster differed transcriptionally between WT and P23H. Transcripts of certain genes involved in the cAMP response, including *junba*, *junbb* and *rgs16*, showed increased expression in P23H whereas transcripts involved in the regulation of cytoskeleton structure including *tubb5*, *tuba1c* and *tuba8l4* are highly expressed in the WT dataset (Fig. 13A). The group of chaperonin T-complex protein genes involved in neuronal development and eye morphogenesis [79], including *cct2*, *cct3*, *cct4*, *cct5* and *cct7* is decreased in the P23H NPCs compared to the WT (Fig. 13B). The WT NPCs also express an increased amount of certain immunoproteasome subunits including *psma1*, *psmb1*, *psmb2*, *psmb3*, *psmb6* and *psmb7* (Fig. 13B). It has been shown that proteasomes are essential in maintaining the self-renewal of neural progenitors and the proteasomes decrease with aging [80]. The P23H degeneration condition may mimic aging due to chronic stress and hence we notice a decrease in the chaperone and proteasome DEGs.

**Figure 13.**
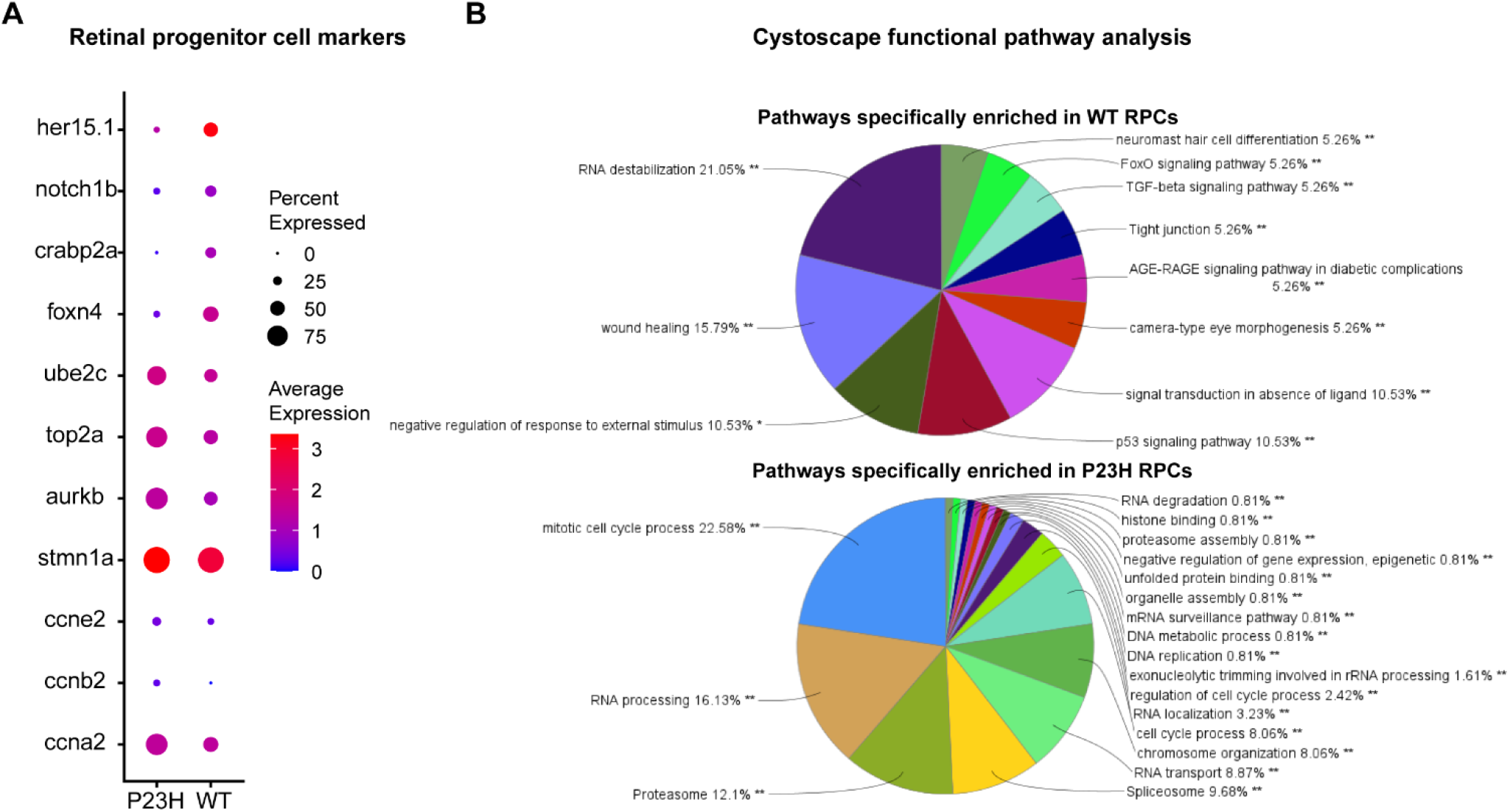
A neurogenic progenitor cell (NPCs) cluster was identified in both WT and P23H. A) Violin plot showing changes in DEGs involved in cAMP response and cytoskeleton structure between WT and P23H PRCs. Dot plot showing the changes in the expression level and the percentage of cells of DEGs involved chaperonin pathway and immunoproteasome system between WT and P23H PRCs.

Overall this study provides a comprehensive transcriptome analysis of most of the predominant retinal cell types of the Zebrafish retina and a first-time comprehensive analysis of the transcriptome changes happening in every retinal cell type during RP in a regeneration-enabled Zebrafish model.

## Discussion

The development of precise molecular and cellular strategies for regeneration-based therapies for RP requires a comprehensive understanding of the cell type-specific changes and cellular heterogeneity that comprise retina remodeling during RP. Interventions that presume substantial preservation of the neural retina will likely fail in the late stages of the disease. Accordingly, our single-cell transcriptome analysis identified molecular pathways that are dysregulated in specific retinal cell types in an RP model, allowing us to create an extensive atlas of cellular changes happening during RP. We found that chronic degeneration and regeneration of rods affect all retinal cell types. Transcriptional changes were diverse, reflecting widespread oxidative stress pathways, modifications of metabolism, changes in circadian rhythm, glutamate signaling and synapse remodeling.

Rod photoreceptor degeneration reduces oxygen consumption in the outer retina and leads to increased oxygen tension that can have damaging consequences for other cells in the outer retina [81]. A disparity between ROS production and antioxidant capacity leads to oxidative damage, which has been reported in many previous studies [6, 54, 82]. Both mature rods and newly-forming rods in the P23H retina display increased levels of oxidative stress markers. We also see a notable increase of genes that deal with oxidative stress such as *sod1*, *sod2* and *hsf2* in cones and RPE, and to a lesser extent in bipolar and amacrine cells, revealing an extensive response to the increase in ROS production.

The retina is one of the most energy-demanding tissues in the body, with very high demand driven by photoreceptors. Despite rod photoreceptor loss in the P23H retina, we see a host of metabolic changes throughout the retina that suggest adaptations to increased energy demand. Glycolysis and TCA cycle enzymes are increased in both rods and cones. While in rods this could be due to increased energy demand to deal with misfolded protein, the notable downregulation of Rod-derived Cone Viability Factor *nxnl1* in rods and cones suggests that glucose uptake could be compromised [29], leading to nutrient starvation and potentially compensatory upregulation of glycolytic and TCA cycle genes. In the inner retina, retinal ganglion cells displayed increased expression of mitochondrial electron transport chain genes, which may reflect energy demand derived from elevated spontaneous activity.

Aside from ATP generation, glycolysis intermediates are the substrates for glycan synthesis, pentose phosphate pathway (PPP), and serine synthesis pathway. We have noticed that genes related to PPP are increased in the P23H cones compared to WT. A recent study by Sinclair et al. [83] showed that metabolic reprogramming is required for successful tail regeneration and an obligate metabolic shift to promote glucose metabolism is necessary for regeneration. Similarly, another study shows that stimulation of glycolysis by enhanced expression of enzymes involved in regulating glucose and pyruvate metabolism promotes cardiomyocyte proliferation and regeneration after injury in adult Zebrafish [84]. Combined with our results, these observations suggest that a metabolic shift towards glycolysis is pronounced during regeneration and needs to be considered for regeneration therapy.

The retina has an endogenous circadian clock that regulates many physiological functions within the retina including POS shedding and phagocytosis, retinomotor movements and synaptic ribbon changes. Dampening and desynchronization of circadian rhythms from zeitgebers were seen in a rat model of RP when photoreceptor degeneration was advanced [85]. Surprisingly, we see an elevated expression of certain transcripts involved in regulating the circadian rhythm in the P23H Zebrafish, including *per1b*, *per2*, *cry1aa* and *tefa*, in the cones, bipolar cells and RPE. Most of these genes are transcriptional repressors and are generally expressed in the same phase of the circadian cycle [46]. The WT and P23H single-cell transcriptome libraries we used were prepared at the same time of day from animals with the same environmental light cycle. There may be a phase shift in the circadian rhythm in P23H, which needs to be further investigated. Some of the elevated circadian genes are also components of the *hsf2* stress response pathway. It is possible that responses to stress in the cell types physically close to the degenerating rods drive broader changes in the circadian regulators. In the present model, we also encounter continuous photoreceptor regeneration and hence the changes may not be comparable to RP models with advanced degeneration. At late stages the circadian rhythm might be further modified, suggesting a cascade of circadian controlled activities in the retina are affected. It has been shown in many studies that the disruption of the circadian pattern is associated with many pathological conditions including obesity, cancer and blindness [86–89]. In this case, their circadian rhythms would be expected to exhibit a non-24-hr pattern. A recent study has shown that melatonin supplementation decreased the rate of desynchronization of the circadian rhythm and improved visual function in P23H rats [90]. This provides a new possibility for treating RP through circadian synchronization and hence it is important to understand the changes in the circadian pattern in different models of RP to get a better understanding of the role of circadian rhythm on RP.

Our data show evidence for significant synapse and axon remodeling throughout the retina in the P23H Zebrafish. In the rods and cones, genes involved in the regulation of neurotransmitter levels, neurotransmitter release, and regulation of synaptic plasticity are upregulated in the P23H retina. In contrast, genes involved in axon guidance and axon extension are reduced. These changes may be compensatory mechanisms in the metabolically and physiologically stressed photoreceptors to maintain some synaptic transmission. Following photoreceptors, horizontal and bipolar cells in P23H show increases in genes involved in axon guidance and dendritic self-avoidance, respectively, suggesting active synaptic remodeling. Unlike mammalian retinal degeneration models, degeneration/regeneration models such as Zebrafish need to accommodate synapse formation with the newly forming rods, making them a highly dynamic system in which to understand synaptic remodeling.

In the inner retina, we noted increased expression of genes involved in glutamate signaling, especially certain ionotropic glutamate receptors, in some amacrine and ganglion cell clusters of P23H compared to WT. This correlates well with enhanced spontaneous activity and oscillatory behaviors in the retina of mammalian models of retinal degeneration [16, 91] and specifically with signatures of enhanced ionotropic glutamatergic input to amacrine and ganglion cells in retinal degeneration [12]. This suggests that despite the ongoing regeneration of rods, the Zebrafish model of RP displays features very similar to mammalian RP models.

Neuronal remodeling during retinal degenerative diseases stands as one of the greatest barriers to implementing regenerative or bionic therapies to restore vision [13, 92]. Our study provides a foundation to understand the molecular underpinnings of the neuronal remodeling that occurs and will help us to understand the steps that need to be taken for successful vision restoration.

## Methods and Materials

### Animal Husbandry

Rearing, breeding, and staging of Zebrafish (*Danio rerio)* were performed following standard procedures in the Zebrafish community [93]. Wild type AB Zebrafish were purchased from the Zebrafish International Resource Center (ZIRC; Eugene, OR, USA; RRID: ZIRC_ZL1), raised, bred, and maintained on a 14 h light/10 h dark cycle. P23H rhodopsin transgenic Zebrafish (ZFIN name uth4Tg; ID ZDB-ALT-210330-3) were developed in-house and characterized in a previous publication [24]. Randomly selected fish between 6 and 10 months of age were used for all experiments. All procedures employing animals comply with the U.S. Public Health Service policy on humane care and use of laboratory animals and the NRC Guide for the Care and Use of Laboratory Animals and have been reviewed and approved by the Institutional Animal Care and Use Committee at the University of Texas Health Science Center at Houston under protocols HSC-AWC-15-0057 and HSC-AWC-18-0047.

### Zebrafish retinal dissociation and single-cell cDNA library preparation

For each sample submitted for single-cell RNA sequencing (sc-RNA Seq), 3 fish of the same strain were collected in the morning between nine and eleven a.m. in Falcon tubes containing 0.15% Tricaine/MS222 and incubated on ice for 10 minutes. Zf were decapitated and eyeballs were detached and placed in dissecting medium containing Leibovitz’s L-15 medium (Gibco 21083027). Each retina was then extracted and placed in a 1.5 mL microcentrifuge tube with 300 μL of dissociation solution containing L15 and EBSS (Gibco 14155063) in a 3:1 ratio. After all retinas were collected, the dissociation solution was replaced with 200 μL of Papain solution (50 U/mL; Worthington Biochemical LK003178) and retinas were incubated at 28°C. After 20 minutes of incubation, retinas were gently triturated 20-30 times with fire-polished glass pipettes. Supernatants were collected and incubated with Papain for another 40 minutes with trituration performed every 10 minutes. Papain reaction was terminated by adding 2x volume of 0.1% BSA in L15: EBSS medium. Tubes were centrifuged at 400g for 5 minutes to pellet the cells. The supernatant was removed and washed again with 0.05% BSA. The supernatant was removed and cells were finally suspended in 100 μl of 0.05% BSA. Cells were checked for cell viability and counted to get a final concentration of at least ~800 to 1000 cells/μl.

The cell samples were submitted for 3’ sc-RNA library preparation and sequencing through the Baylor College of Medicine Single Cell Genomics Core facility. Single-cell Gene Expression Libraries were prepared using Chromium Next GEM Single Cell 3’ Reagent Kits v2 and v3.1 (10x Genomics, Pleasanton, CA). In brief, single cells, reverse transcription (RT) reagents, Gel Beads containing barcoded oligonucleotides, and oil were loaded on a Chromium controller (10x Genomics) to generate single cell GEMS (Gel Beads-In-Emulsions) where full-length cDNA was synthesized and barcoded for each single cell. Subsequently, the GEMS are broken and cDNA from every single cell is pooled. Following cleanup using Dynabeads MyOne Silane Beads, cDNA is amplified by PCR. The amplified product is fragmented to optimal size before end-repair, A-tailing, and adaptor ligation. The final library was generated by amplification. cDNA libraries were sequenced on an Illumina HiSeq2500.

Sequences were run through the 10x Genomics CellRanger V3.1.2.0 pipeline. First, sequences were demultiplexed into FASTQ files through CellRanger mkfastq. FASTQ files were then aligned to the Zf reference genome (GCA_000002035.4_GRCz11) through CellRanger count. Output files were run through Babraham Bioinformatics’ FASTQC V0.11.9 to check on the quality of the sequences used for analysis. Sequences had a normal distribution of GC content as well as base call accuracy above 99%. Over 93% of reads mapped successfully to the genome. All computational analysis was performed through Texas Advanced Computing Center (TACC) Lonestar5 computing service. Scripts were written in Notepad++ v7.6.2 and uploaded to TACC through PuTTY V0.72.

### Single-cell RNA-seq data analysis

The total number of DNA sequence reads for v3.1 datasets were 608,597,236 for P23H and 967,526,548 for WT. All reads were aligned to the Zf reference genome (GCF_000002035.6_GRCz11) through Texas Advanced Computing Center’s (TACC’s) Lonestar5 computing system. After alignment, quality control was assessed through FASTQC. Aligned data were run through the Seurat V3.1.1 package [25] in R V3.6.1. Any cell that was determined to be an outlier or contained greater than 10% mitochondrial genes was removed. Cells initially reported for analysis were 16,516 for P23H and 16,979 for WT datasets. From a total of 16,979 cells (P23H), we obtained an average of 35,844 reads per cell and 1,514 UMIs (unique transcripts) per cell. We detected a total of 22,497 genes from the P23H dataset. From a total of 16,516 cells (WT), we obtained an average of 58,581 reads per cell and 1,535 UMIs per cell. We detected a total of 21,889 genes from the WT dataset. The reported median genes for P23H and WT datasets were 665 and 610 genes, respectively. P23H and WT datasets contained between 200 and 4,700 genes and 200 and 6,000 genes per cell, respectively (Table S4). Low abundant genes (expressed in less than 10 cells) and cells of potentially low quality (percentage of mitochondrial genes >10%) were removed from downstream analysis. PCElbowPlots were performed and 20 principal components were used for downstream analysis of each dataset. PC1 to PC20 were used to construct nearest neighbor graphs in the PCA space followed by Louvain clustering and non-linear dimensional reduction by tSNE to visualize and explore the clusters. Each cluster was assigned a cell type based on their expression of cell type-specific marker genes (Table S1). Dot plots, Heatmaps, Violin Plots, and tSNE cluster maps were all generated through Seurat’s vignettes V3.2. Expression levels are expressed in a base 2 log scale. Statistical significance between expression levels was determined using Wilcoxon ‘Mann-Whitney’ test (two-sided, unpaired with 95% confidence level).

### Network Analysis

Differentially expressed genes (DEGs) for each cluster were uploaded to the PANTHER classification system V16.0 [94, 95] (http://pantherdb.org/) to identify the different functional classifications groups expressed. DEGs for each cluster were then imported into Cytoscape’s plugin ClueGo 3.9.0 [27] (https://cytoscape.org/) to visualize the non-redundant biological networks involved for each cluster. Pathways that show a pValue less than 0.05 were used for analysis. Networks were assessed to determine the functional roles and differences between the P23H and WT datasets. We also used the tool Metascape [28] (https://metascape.org/) for a comprehensive analysis of the differentially expressed genes (DEGs) from different cell types. Functional Enrichment Analysis to determine protein-protein interaction networks was performed on genes of interest through STRING V11.0 (https://string-db.org/).

### Fluorescent In-Situ Hybridization Multiplex/HiPlex

All fish for tissue analysis were collected during the morning between nine and eleven a.m. and anesthetized in 0.15% of Tricaine/MS222 on ice. Fish were then decapitated and eyeballs enucleated and fixed in 4% formaldehyde (PFA) in 0.1 M phosphate buffer (PB), pH 7.5 for 24hr at RT. Eyes were then washed four times at 15-minute intervals in PB and cryoprotected in 30% sucrose in PB overnight at 4°C. Eyes were then frozen in Tissue-Tek O.C.T. compound (Sakura Finetek, Torrance, CA, USA) using dry ice and stored at −80°C. Retinal sections were cut at 20 μm on a cryostat (Leica) and stored at −80°C until use. We used the RNAscope Hiplex detection reagents supplied by the manufacturer (Advanced Cell Diagnostics, Newark, CA, USA) along with the specific probes for target genes. HiPlex and subsequent fluorescent multiplex assays were performed following the manufacturer’s instructions [96]. Briefly, sections were fixed in 4% PFA, dehydrated with 50%, 70%, and 100% ethanol, then treated with protease (from the HiPlex kit 324419). In HiPlex, all gene targets are hybridized and amplified together, whereas detection is achieved iteratively in groups of 2-3 targets. Sections were incubated with pooled HiPlex probes and amplified with a series of amplification solutions, then a detection solution was applied. Samples were hybridized with probes, incubated with amplification solution, signals were visualized by applying detection solution. Cell nuclei were counterstained with DAPI and samples were mounted. Images were taken using a Zeiss LSM 800 laser scanning confocal microscope. After each round, samples were treated with TCEP solution to cleave fluorophores (from the HiPlex kit 324419), then moved on to the next round of the fluorophore detection procedures.

## Acknowledgments

This work was supported by a grant from the William Stamps Farish Fund (JOB), the Louisa Stude Sarofim endowment (JOB), facility grant G20RR024000, core grants P30EY028102 and P30EY007551, and the University of Houston College of Optometry. Single-cell library preparation and sequencing at the Baylor College of Medicine Single Cell Genomics Core was partially supported by NIH shared instrument grants S10OD023469 and S10OD025240, core grant P30EY002520 and CPRIT grant RP200504.

