## Supplemental Figures and Tables for "Transcriptomic remodeling of the retina in a Zebrafish model of Retinitis Pigmentosa"

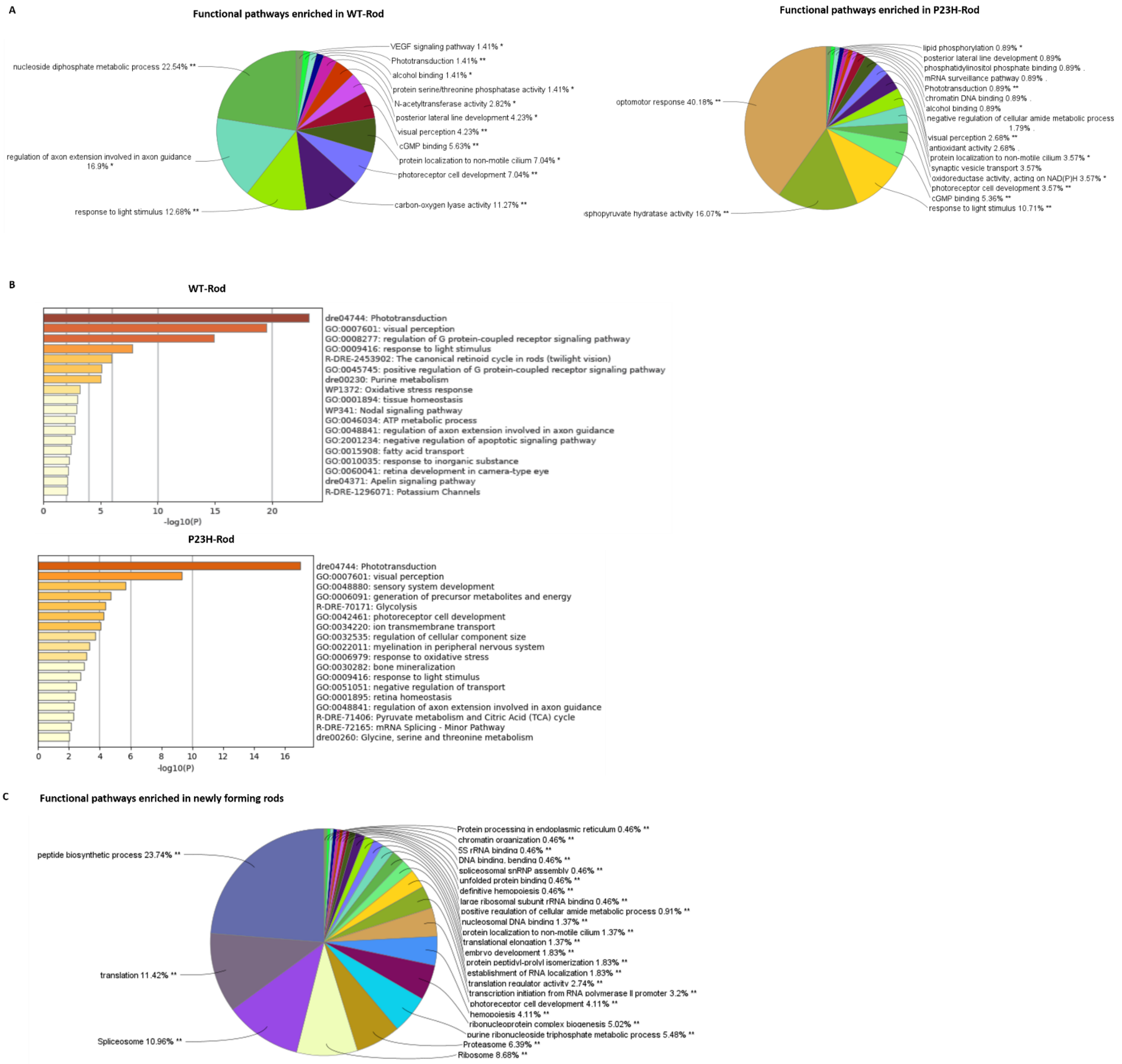


Figure S1. **Changes associated with rod functional pathways between WT and P23H.** A) Functional pathways specifically present in the WT and P23H rod cells are revealed by Cytoscape analysis as described in the methods. B) Functional pathways specifically present in the WT and P23H rod cells revealed by Metascape analysis. C) Functional pathways specifically enriched in the newly formed rod cells revealed by Cytoscape analysis.

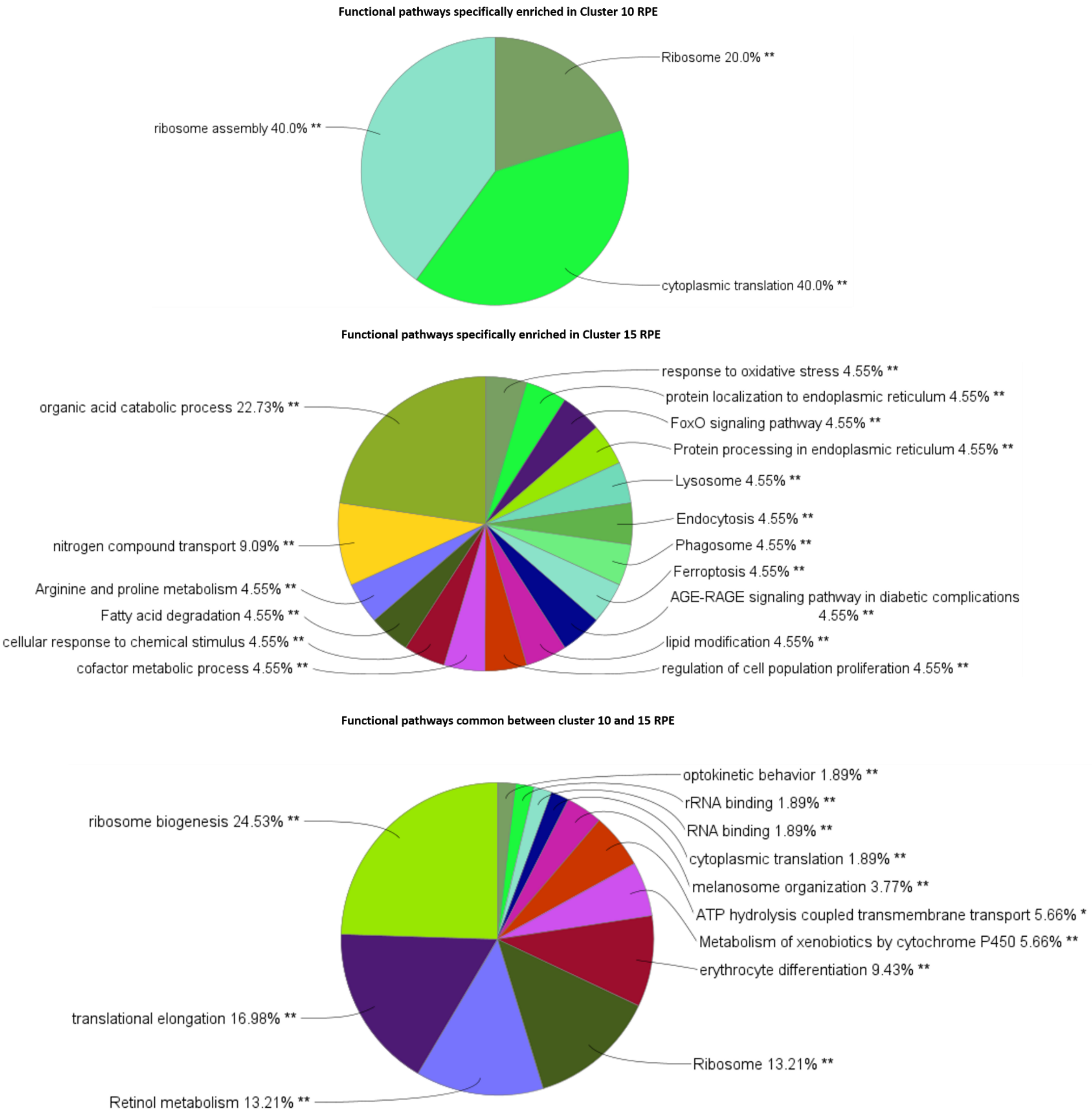


Figure S2. **Functional pathway analysis on RPE cells by Cytoscape.** Functional pathways specifically present commonly between 10 and 15 RPE clusters as well as pathways specifically enriched in clusters 10 and 15.

A

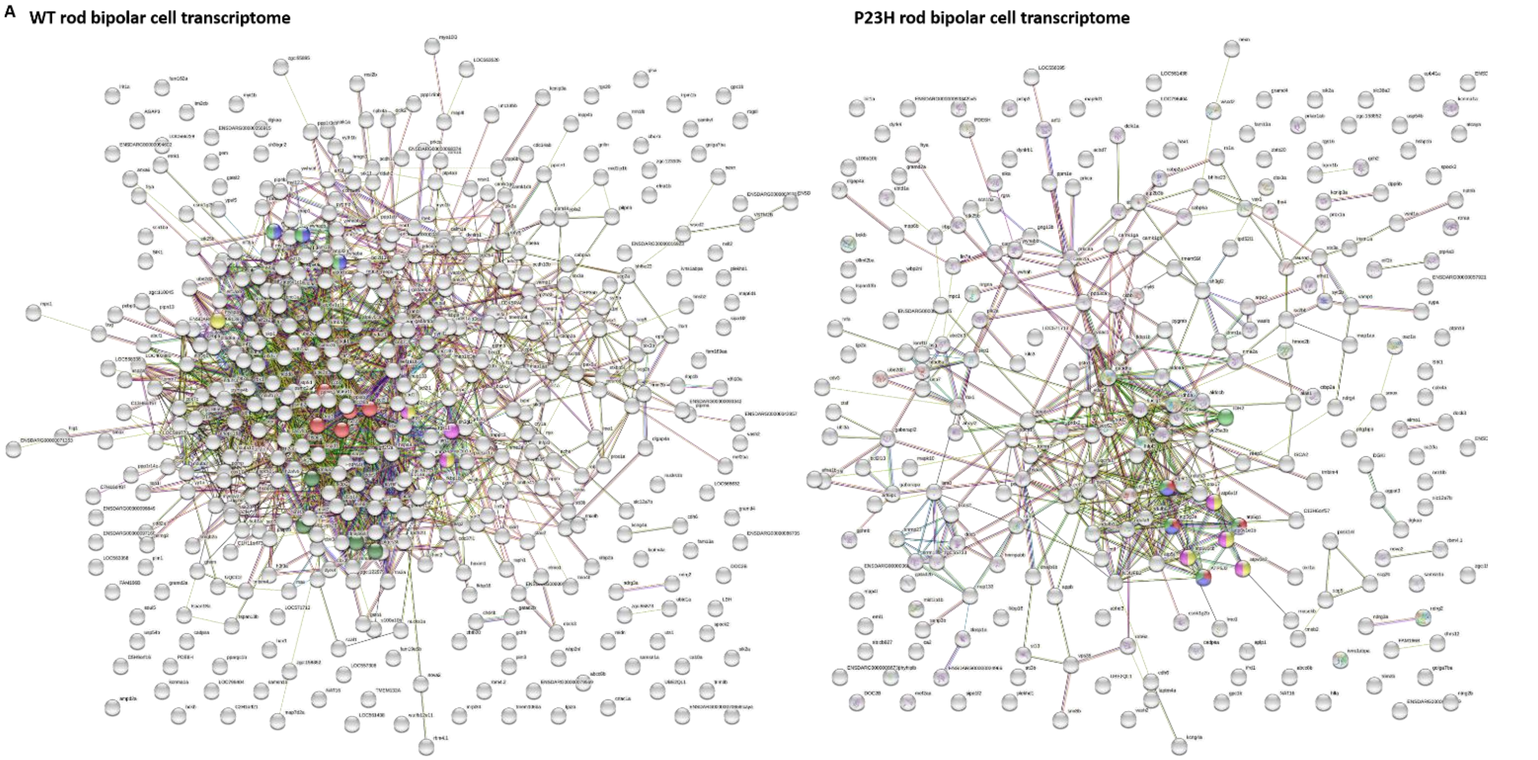


B

Reactome Pathways

| pathway | description | count in network | strength | false discovery rate |
|---|---|---|---|---|
| DRE-390471 | Association of TriC/CCT with target proteins during biosynt... | 6 of 9 | 1.81 | 2.80e-06 |
| DRE-8866427 | VLDLR internalisation and degradation | 3 of 11 | 1.42 | 0.0071 |
| DRE-111447 | Activation of BAD and translocation to mitochondria | 2 of 9 | 1.33 | 0.0280 |
| DRE-1445148 | Translocation of SLC2A4 (GLUT4) to the plasma membrane | 2 of 10 | 1.29 | 0.0312 |
| DRE-3371497 | HSP90 chaperone cycle for steroid hormone receptors (SHR) | 4 of 21 | 1.27 | 0.0043 |

Reactome Pathways

| pathway | description | count in network | strength | false discovery rate |
|---|---|---|---|---|
| DRE-8949613 | Cristae formation | 4 of 20 | 1.31 | 0.0046 |
| DRE-163210 | Formation of ATP by chemiosmotic coupling | 4 of 20 | 1.31 | 0.0046 |
| DRE-1592230 | Mitochondrial biogenesis | 5 of 32 | 1.2 | 0.0026 |
| DRE-77387 | Insulin receptor recycling | 4 of 29 | 1.15 | 0.0102 |
| DRE-1222556 | ROS, RNS production in phagocytes | 4 of 31 | 1.12 | 0.0119 |

Figure S3. **Changes associated with rod bipolar cell functional pathways between WT and P23H.** A) Changes in the transcriptome interaction network between WT and P23H were revealed by STRING interaction analysis. B) Reactome pathways enriched in WT vs P23H rod bipolar cells by STRING analysis.

Functional pathways specifically enriched in WT-HC

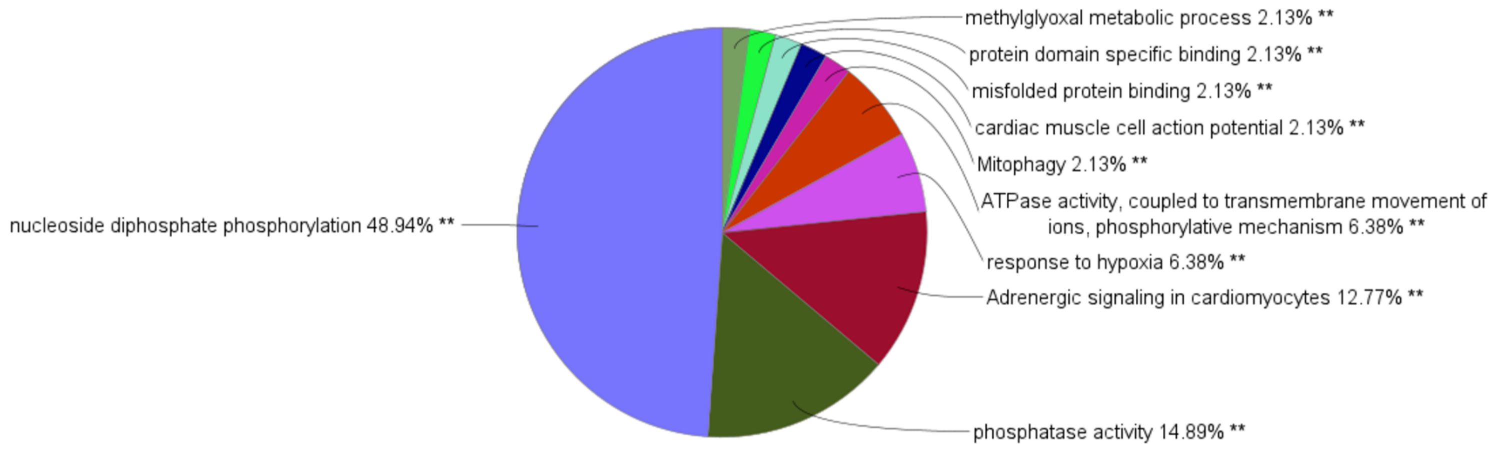


Functional pathways specifically enriched in P23H-HC

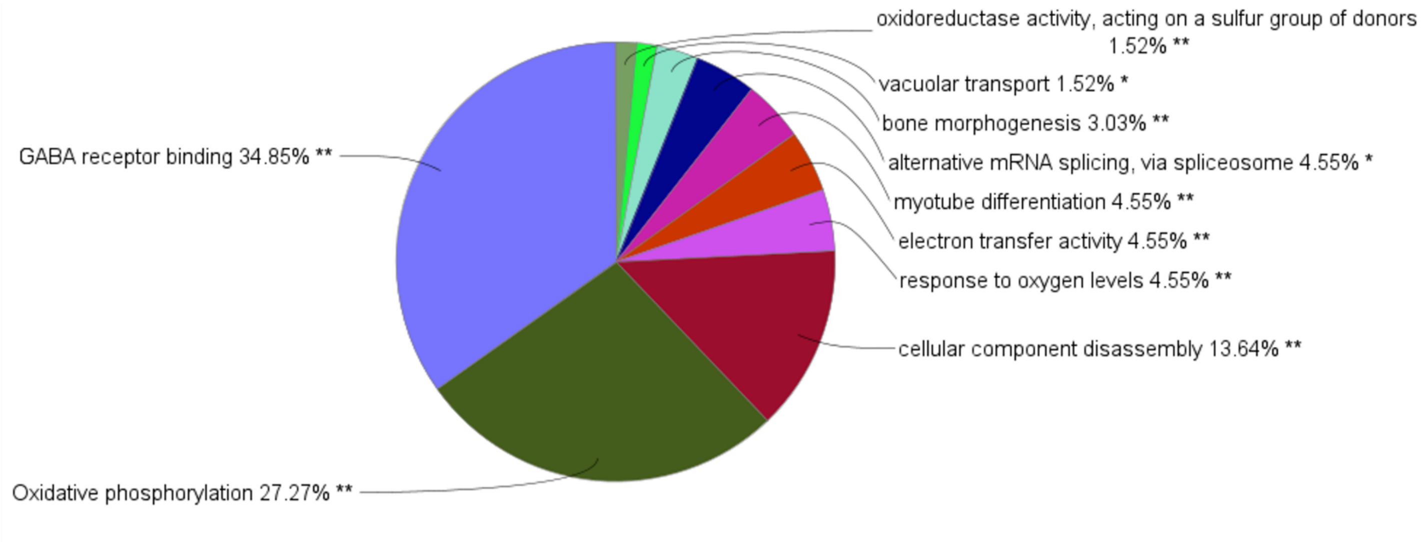


Figure S4. **Functional pathway analysis on Horizontal cells by Cytoscape.** Functional pathways specifically present in the WT and P23H Horizontal cells.

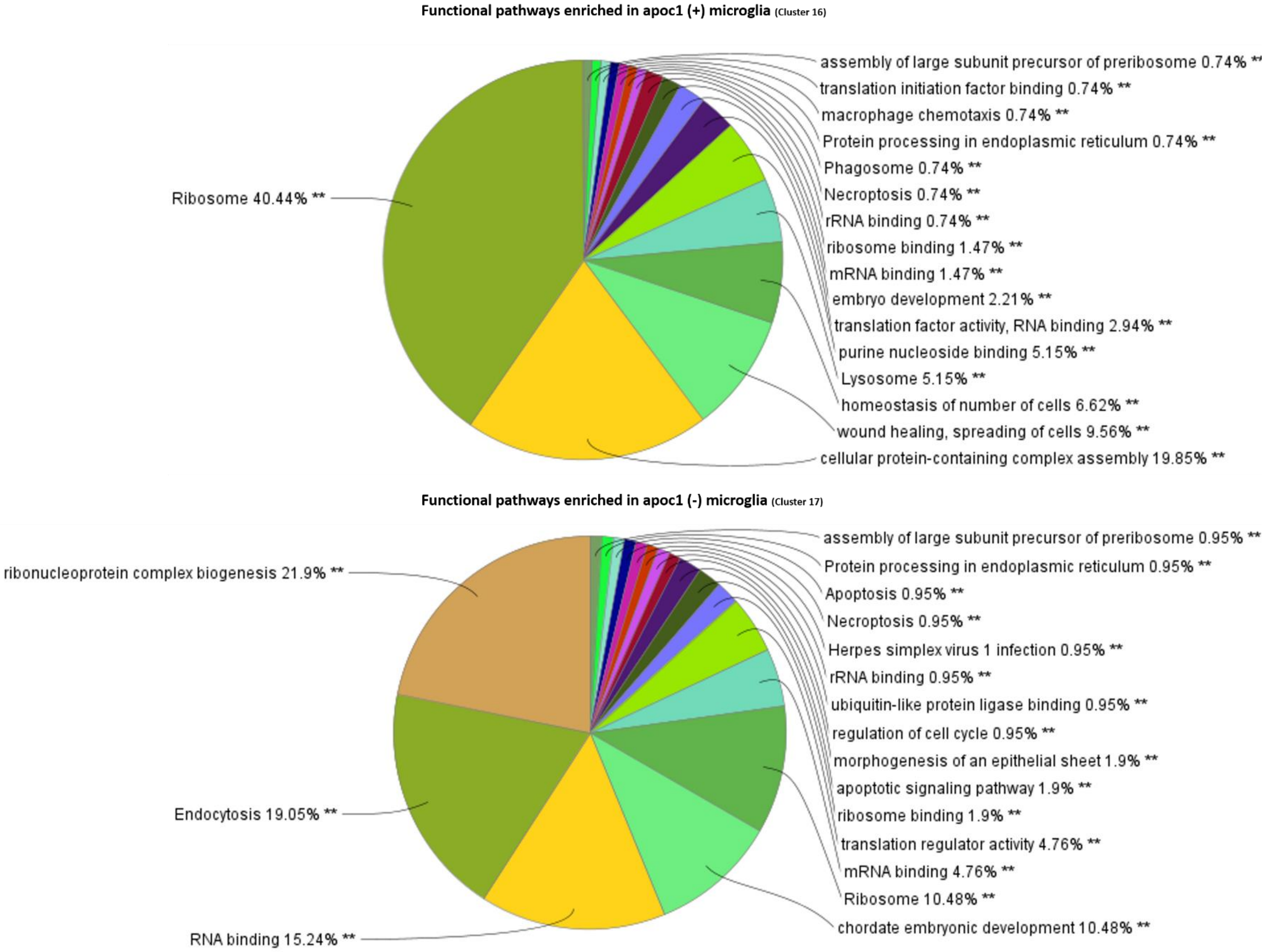


Figure S5. **Functional pathway analysis on microglia/macrophages by Cytoscape.** Functional pathways specifically enriched in the apoc1(+) and apoc1(-) microglia/macrophages.

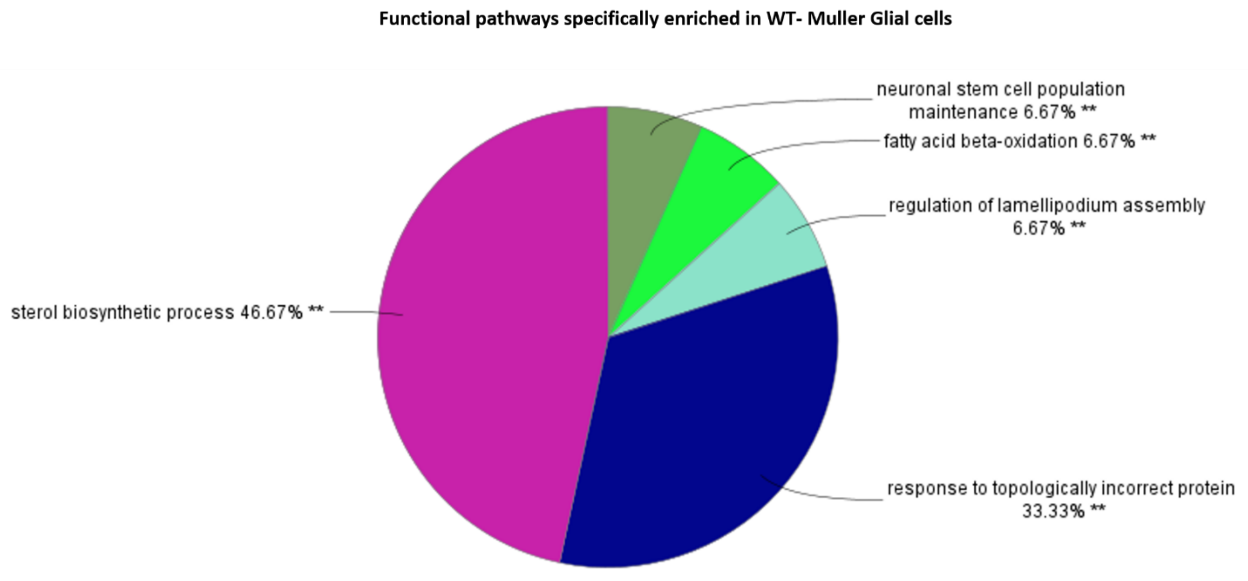


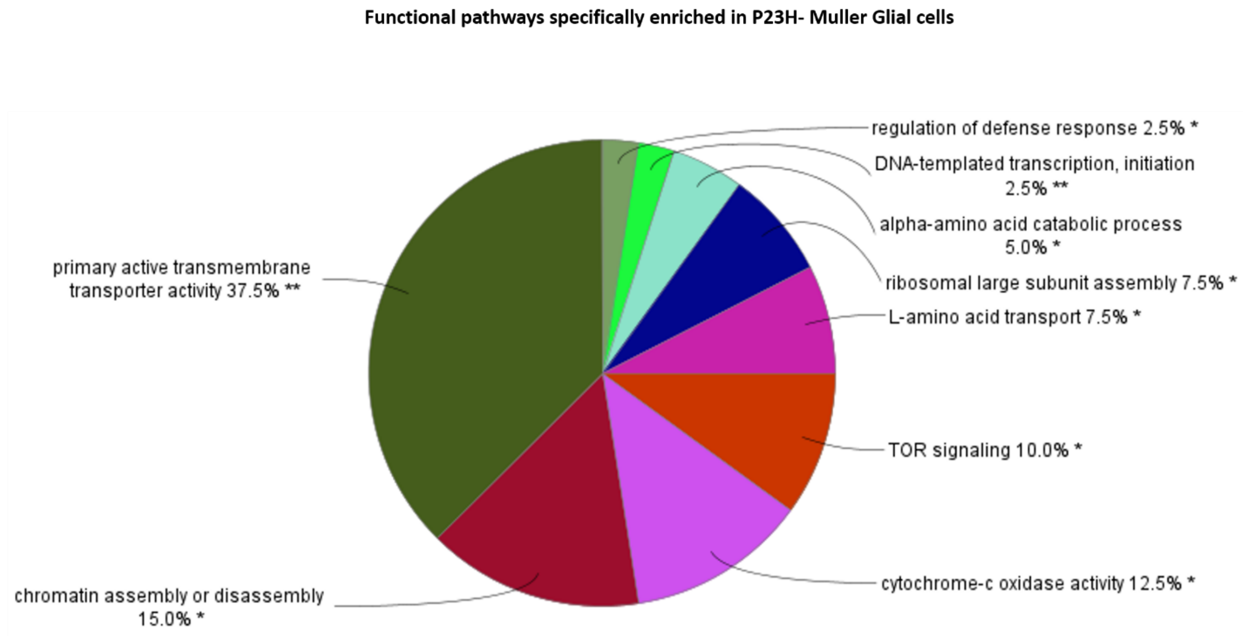


Figure S6. **Functional pathway analysis on Müller glial cells by Cytoscape.** Functional pathways specifically present in the WT and P23H Müller glial cells.

Table S1. **List of marker genes used to identify cell types with references**

| Gene markers | Cell Type | Gene ID | Source |
|---|---|---|---|
| cabp1a | Amacrine Cells | 449789 | Characterization of the Calcium Binding Protein Family in Zebrafish [1] |
| tfap2a | Amacrine Cells | 140618 | Tfap2a and 2b act downstream of Ptf1a to promote amacrine cell differentiation during retinogenesis [2] |
| syt2a | Amacrine Cells | 567877 | Localization of the Calcium-binding Protein Secretagogin in Cone Bipolar Cells of the Mammalian Retina [3] |
| chata, gad2 | Amacrine Cells | 100170938, 550403 | Molecular identification of sixty-three amacrine cell types completes a mouse retinal cell atlas [4] |
| cabp2a | Bipolar Cells | 572226 | Characterization of the Calcium Binding Protein Family in Zebrafish [1] |
| prkca | Bipolar Cells | 497384 | A comparative analysis of rod bipolar cell transcriptomes identifies novel genes implicated in night vision [5] |
| vsx1 | Bipolar Cells | 30598 | Vsx2 in the zebrafish retina: restricted lineages through derepression [6] |
| rs1a | Bipolar Cells | 445044 | Identification of molecular markers of bipolar cells in the murine retina [7] |
| bhlhe23 | Bipolar Cells | 559796 | A comparative analysis of rod bipolar cell transcriptomes identifies novel genes implicated in night vision [5] |
| gnat1 | Rods | 140428 | Transcripts within rod photoreceptors of Zebrafish Retina [8] |
| rom1a, rom1b | Rods | 393767, 393989 | Rom-1 is required for rod photoreceptor viability and the regulation of disk morphogenesis [9] |
| saga, sagb | Rods | 792319, 619268 | Transcripts within rod photoreceptors of Zebrafish Retina [8] |
| gnat2 | Cones | 140429 | Transducin Duplicates in the Zebrafish Retina and Pineal Complex: Differential Specialisation after the Teleost Tetraploidisation [10] |
| pde6c | Cones | 393845 | A Mutation in the Cone-Specific pde6 Gene Causes Rapid Cone Photoreceptor Degeneration in Zebrafish [11] |
| clul1 | Cones | 559018 | Comparative Analysis and Expression of CLUL1, a Cone Photoreceptor-Specific Gene [12] |
| cx52.6 | Horizontal Cells | 404207 | Specific connectivity between photoreceptors and horizontal cells in the zebrafish retina [13] |
| cx55.5 | Horizontal Cells | 573483 | Specific connectivity between photoreceptors and horizontal cells in the zebrafish retina [13] |
| cx52.9 | Horizontal Cells | 404625 | Specific connectivity between photoreceptors and horizontal cells in the zebrafish retina [13] |
| her4.3 | Retinal Progenitor Cells | 792198 | Tracking the fate of her4 expressing cells in the regenerating retina using her4:Kaede zebrafish [14] |
| insm1a | Retinal Progenitor Cells | 402941 | Insm1a-mediated gene repression is essential for the formation and differentiation of Müller glia-derived progenitors in the injured retina [15] |

| | | | |
|---|---|---|---|
| her15.1 | Retinal Progenitor Cells | 100534909 | The transcription factor hairy/E(spl)-related 2 induces proliferation of neural progenitors and regulates neurogenesis and gliogenesis [16] |
| rpe65a | Retinal Pigment Epithelial Cells | 393724 | Expression profiling of the RPE in zebrafish smarca4 mutant revealed altered signals that potentially affect RPE and retinal differentiation [17] |
| dct | Retinal Pigment Epithelial Cells | 58074 | Expression profiling of the RPE in zebrafish smarca4 mutant revealed altered signals that potentially affect RPE and retinal differentiation [17] |
| pmela, pmelb | Retinal Pigment Epithelial Cells | 321239, 562810 | Identification of cell surface markers and establishment of monolayer differentiation to retinal pigment epithelial cells [18] |
| rbpms2a, rbpms2b | Retinal Ganglion Cell | 436682 | The RNA binding protein RBPMS is a selective marker of ganglion cells in the mammalian retina [19] |
| pou4f1 | Retinal Ganglion Cell | 58057 | Genetic interplay between transcription factor Pou4f1/Brn3a and neurotrophin receptor Ret in retinal ganglion cell type specification [20] |
| isl2b | Retinal Ganglion Cell | 30151 | Molecular classification of zebrafish retinal ganglion cells links genes to cell types to behavior [21] |
| plp1b | Oligodendrocytes | 368234 | Identification of genes expressed by zebrafish oligodendrocytes using a differential microarray screen [22] |
| cd59 | Oligodendrocytes | 567192 | Cd59 and inflammation regulate Schwann cell development [23] |
| cd9b | Oligodendrocytes | 406737 | Expression and distribution of CD9 in myelin of the central and peripheral nervous systems [24] |
| gfap | Muller Glial Cells | 30646 | Zebrafish Lbh-like Is Required for Otx2-mediated Photoreceptor Differentiation [25] |
| cahz | Muller Glial Cells | 30331 | Characterization of Müller glia and neuronal progenitors during adult zebrafish retinal regeneration [26] |
| ptgdsb.1 | Muller Glial Cells | 336492 | Rapid, Dynamic Activation of Müller Glial Stem Cell Responses in Zebrafish [27] |
| cd74a | Microglia | 58113 | Regeneration associated transcriptional signature of retinal microglia and macrophages [28] |
| cd74b | Microglia | 30645 | Regeneration associated transcriptional signature of retinal microglia and macrophages [28] |
| apoc1 | Microglia | 570638 | Regeneration associated transcriptional signature of retinal microglia and macrophages [28] |

Table S2. **List of Top 10 genes from each cluster (WT)**

| gene | p_val | avg_log2FC | pct.1 | pct.2 | p_val_adj | cluster |
|---|---|---|---|---|---|---|
| cabp5a | 0 | 3.930352 | 0.958 | 0.455 | 0 | 0 |
| cabp2a | 0 | 3.328842 | 0.795 | 0.156 | 0 | 0 |
| rs1a | 0 | 2.985809 | 0.652 | 0.189 | 0 | 0 |
| lrit1a | 0 | 2.933814 | 0.634 | 0.111 | 0 | 0 |
| si:ch73-256g18.2 | 0 | 2.905666 | 0.837 | 0.17 | 0 | 0 |
| vamp1 | 0 | 2.784025 | 0.894 | 0.229 | 0 | 0 |
| efna1b | 0 | 2.776288 | 0.84 | 0.328 | 0 | 0 |
| waslb | 0 | 2.503441 | 0.643 | 0.178 | 0 | 0 |
| vsx1 | 0 | 2.491107 | 0.497 | 0.082 | 0 | 0 |
| nrn1lb | 0 | 2.338138 | 0.524 | 0.088 | 0 | 0 |
| opn1lw2 | 1.22E-66 | 4.441128 | 0.286 | 0.11 | 2.3E-62 | 1 |
| opn1lw1-1 | 3.9E-186 | 4.40282 | 0.447 | 0.133 | 7.4E-182 | 1 |
| opn1lw2-1 | 4.9E-113 | 4.352769 | 0.558 | 0.289 | 9.2E-109 | 1 |
| opn1mw2 | 2.87E-37 | 3.886074 | 0.265 | 0.135 | 5.42E-33 | 1 |
| clul1 | 0 | 3.494121 | 0.938 | 0.282 | 0 | 1 |
| hexb | 0 | 3.365444 | 0.806 | 0.179 | 0 | 1 |
| LOC100334711 | 0 | 3.099652 | 0.768 | 0.112 | 0 | 1 |
| prph2a | 0 | 3.064058 | 0.892 | 0.204 | 0 | 1 |
| arr3a-1 | 2.3E-269 | 2.866967 | 0.868 | 0.555 | 4.3E-265 | 1 |
| htra1b | 0 | 2.860907 | 0.823 | 0.13 | 0 | 1 |
| gad2 | 0 | 3.287965 | 0.926 | 0.128 | 0 | 2 |
| LOC557301 | 1.2E-120 | 3.127811 | 0.335 | 0.093 | 2.3E-116 | 2 |
| crhbp | 1.27E-60 | 2.722586 | 0.254 | 0.09 | 2.4E-56 | 2 |
| scg2b | 0 | 2.641145 | 0.676 | 0.108 | 0 | 2 |
| egr4 | 5.3E-213 | 2.624786 | 0.689 | 0.247 | 1E-208 | 2 |
| slc6a1b | 0 | 2.58459 | 0.855 | 0.08 | 0 | 2 |
| snap25a | 0 | 2.576376 | 0.957 | 0.329 | 0 | 2 |
| slc6a1a | 0 | 2.56403 | 0.865 | 0.067 | 0 | 2 |
| vamp2 | 0 | 2.453991 | 0.959 | 0.294 | 0 | 2 |
| stxbp1a | 0 | 2.361369 | 0.895 | 0.155 | 0 | 2 |
| rho | 0 | 3.811814 | 1 | 0.999 | 0 | 3 |
| rom1b | 0 | 3.503928 | 0.839 | 0.44 | 0 | 3 |
| si:ch211-113d22.2 | 0 | 3.366633 | 0.88 | 0.607 | 0 | 3 |
| si:dkey-22i16.2 | 0 | 3.30342 | 0.795 | 0.314 | 0 | 3 |
| cnga1 | 0 | 3.213976 | 0.773 | 0.27 | 0 | 3 |
| gnat1 | 0 | 3.16765 | 0.989 | 0.937 | 0 | 3 |
| rhol | 0 | 2.969203 | 0.657 | 0.144 | 0 | 3 |
| pde6g | 9E-276 | 2.940326 | 0.91 | 0.758 | 1.7E-271 | 3 |
| rom1a | 0 | 2.721132 | 0.645 | 0.169 | 0 | 3 |

| | | | | | | |
|---|---|---|---|---|---|---|
| elovl4b | 2E-299 | 2.710312 | 0.73 | 0.299 | 3.8E-295 | 3 |
| dct | 0 | 4.388483 | 0.937 | 0.144 | 0 | 4 |
| lrp1aa | 0 | 4.241001 | 0.837 | 0.125 | 0 | 4 |
| pmela | 0 | 4.231526 | 0.87 | 0.087 | 0 | 4 |
| ambp-1 | 0 | 4.227394 | 0.883 | 0.124 | 0 | 4 |
| rgrb | 0 | 4.171871 | 0.99 | 0.453 | 0 | 4 |
| cst3 | 0 | 3.888058 | 0.999 | 0.911 | 0 | 4 |
| pttg1ipb | 0 | 3.876994 | 0.904 | 0.208 | 0 | 4 |
| tyrp1b | 0 | 3.796407 | 0.896 | 0.132 | 0 | 4 |
| mb | 7.1E-138 | 3.639 | 0.484 | 0.189 | 1.3E-133 | 4 |
| itih1 | 0 | 3.627927 | 0.701 | 0.058 | 0 | 4 |
| slc6a1b | 9.7E-165 | 2.822531 | 0.45 | 0.118 | 1.8E-160 | 5 |
| slc6a1a | 1.3E-203 | 2.753738 | 0.464 | 0.105 | 2.5E-199 | 5 |
| slc6a9 | 3.28E-79 | 2.753095 | 0.297 | 0.094 | 6.2E-75 | 5 |
| grin1a | 1.6E-278 | 2.724345 | 0.751 | 0.234 | 3.1E-274 | 5 |
| slc32a1 | 8.5E-289 | 2.692321 | 0.601 | 0.133 | 1.6E-284 | 5 |
| syt2a | 2.8E-175 | 2.672815 | 0.425 | 0.1 | 5.4E-171 | 5 |
| sv2a | 3.6E-282 | 2.600473 | 0.7 | 0.192 | 6.8E-278 | 5 |
| syt1a | 0 | 2.561834 | 0.804 | 0.221 | 0 | 5 |
| LOC103908668 | 5.2E-152 | 2.318628 | 0.515 | 0.172 | 9.8E-148 | 5 |
| gabrb2 | 9.4E-152 | 2.224089 | 0.403 | 0.102 | 1.8E-147 | 5 |
| arr3b | 0 | 4.137515 | 0.981 | 0.459 | 0 | 6 |
| gngt2b | 0 | 3.367986 | 1 | 0.971 | 0 | 6 |
| es1 | 7.5E-308 | 3.227079 | 1 | 0.973 | 1.4E-303 | 6 |
| zgc:73359 | 0 | 2.952109 | 1 | 0.984 | 0 | 6 |
| gnb3b | 0 | 2.918634 | 1 | 0.773 | 0 | 6 |
| si:dkey-97a13.12 | 0 | 2.768871 | 0.993 | 0.537 | 0 | 6 |
| gnat2 | 0 | 2.720752 | 0.998 | 0.825 | 0 | 6 |
| si:dkey-126g1.9 | 0 | 2.703528 | 0.99 | 0.263 | 0 | 6 |
| si:cabz01076231.1 | 0 | 2.525011 | 0.988 | 0.279 | 0 | 6 |
| kera | 0 | 2.510421 | 0.843 | 0.115 | 0 | 6 |
| LOC103909376 | 2.1E-120 | 2.831975 | 0.389 | 0.088 | 4E-116 | 7 |
| si:ch211-222l21.1 | 1.2E-265 | 2.615485 | 0.603 | 0.102 | 2.3E-261 | 7 |
| cxxc5a | 5.8E-266 | 2.249814 | 0.49 | 0.065 | 1.1E-261 | 7 |
| sox4a | 1.9E-181 | 2.189421 | 0.528 | 0.107 | 3.6E-177 | 7 |
| tmsb | 1.4E-263 | 2.146742 | 0.339 | 0.028 | 2.7E-259 | 7 |
| rplp2l | 2.5E-181 | 2.079537 | 0.954 | 0.62 | 4.7E-177 | 7 |
| rpl27a | 3.2E-157 | 2.048143 | 0.782 | 0.335 | 6E-153 | 7 |
| eef1g | 1.4E-157 | 1.9788 | 0.885 | 0.504 | 2.7E-153 | 7 |
| fabp7a | 7.9E-110 | 1.963597 | 0.472 | 0.129 | 1.5E-105 | 7 |
| rps20 | 8.3E-184 | 1.961726 | 0.984 | 0.789 | 1.6E-179 | 7 |

| | | | | | | |
|---|---|---|---|---|---|---|
| mdka | 0 | 6.630412 | 1 | 0.191 | 0 | 8 |
| si:dkey-273o13.3 | 0 | 5.691251 | 0.981 | 0.011 | 0 | 8 |
| rprmb | 0 | 4.765208 | 0.956 | 0.018 | 0 | 8 |
| aqp9a | 0 | 4.698649 | 0.96 | 0.013 | 0 | 8 |
| lbh | 0 | 4.632958 | 0.907 | 0.034 | 0 | 8 |
| atp1b1a | 0 | 4.525271 | 0.996 | 0.153 | 0 | 8 |
| ompa | 0 | 4.482134 | 0.879 | 0.011 | 0 | 8 |
| LOC101882145 | 0 | 4.450836 | 0.911 | 0.026 | 0 | 8 |
| plekhd1 | 0 | 4.391484 | 0.939 | 0.047 | 0 | 8 |
| slc4a5 | 0 | 4.372114 | 0.883 | 0.031 | 0 | 8 |
| rbpms2a | 0 | 4.935184 | 0.941 | 0.047 | 0 | 9 |
| si:dkey-7j14.5 | 0 | 4.121109 | 0.941 | 0.093 | 0 | 9 |
| rbpms2b | 0 | 3.522256 | 0.836 | 0.038 | 0 | 9 |
| wu:fj58g06-1 | 5.9E-150 | 3.466013 | 0.301 | 0.038 | 1.1E-145 | 9 |
| nrgna | 0 | 3.227394 | 0.738 | 0.116 | 0 | 9 |
| cplx2l | 0 | 3.207703 | 0.818 | 0.114 | 0 | 9 |
| isl2b | 0 | 3.078175 | 0.572 | 0.007 | 0 | 9 |
| rab6ba | 0 | 3.028613 | 0.815 | 0.05 | 0 | 9 |
| adcyap1b | 0 | 2.975482 | 0.469 | 0.005 | 0 | 9 |
| inab | 0 | 2.91146 | 0.711 | 0.014 | 0 | 9 |
| ba1 | 2.6E-251 | 7.558887 | 1 | 0.593 | 4.9E-247 | 10 |
| hbaa1 | 2.7E-242 | 7.446047 | 1 | 0.689 | 5.1E-238 | 10 |
| ba1l | 0 | 6.47716 | 1 | 0.123 | 0 | 10 |
| si:ch211-5k11.8 | 0 | 6.421534 | 1 | 0.209 | 0 | 10 |
| si:ch211-250g4.3 | 0 | 5.997521 | 0.978 | 0.008 | 0 | 10 |
| creg1 | 0 | 5.112928 | 0.973 | 0.086 | 0 | 10 |
| hbba2 | 0 | 4.301128 | 0.603 | 0.013 | 0 | 10 |
| hbaa2 | 0 | 3.759552 | 0.427 | 0.011 | 0 | 10 |
| mibp | 0 | 3.053733 | 0.608 | 0.008 | 0 | 10 |
| blvrb | 0 | 2.790517 | 0.597 | 0.066 | 0 | 10 |
| rpe65a | 0 | 4.437179 | 0.99 | 0.163 | 0 | 11 |
| rlbp1b | 3.6E-280 | 4.27623 | 0.99 | 0.267 | 6.8E-276 | 11 |
| thbs1b | 0 | 3.950067 | 0.997 | 0.096 | 0 | 11 |
| f3b | 0 | 3.873123 | 0.926 | 0.057 | 0 | 11 |
| rlbp1b-1 | 2.6E-281 | 3.780893 | 0.993 | 0.254 | 4.9E-277 | 11 |
| pnp4a | 0 | 3.724114 | 0.987 | 0.059 | 0 | 11 |
| fabp11b | 7.7E-270 | 3.695808 | 0.977 | 0.231 | 1.4E-265 | 11 |
| rbp5 | 8.8E-228 | 3.687865 | 0.993 | 0.352 | 1.7E-223 | 11 |
| dusp2 | 0 | 3.670689 | 0.997 | 0.203 | 0 | 11 |
| ca9 | 0 | 3.629561 | 0.987 | 0.057 | 0 | 11 |
| ptgdsb.2 | 0 | 6.069471 | 0.996 | 0.085 | 0 | 12 |

| | | | | | | |
|---|---|---|---|---|---|---|
| apoeb | 0 | 5.747821 | 0.993 | 0.1 | 0 | 12 |
| zgc:153704 | 0 | 5.616926 | 0.978 | 0.054 | 0 | 12 |
| rlbp1a | 0 | 5.247792 | 1 | 0.061 | 0 | 12 |
| igfbp1a | 0 | 5.134332 | 0.859 | 0.028 | 0 | 12 |
| cebpd | 0 | 4.965995 | 0.909 | 0.02 | 0 | 12 |
| ptgdsb.1 | 0 | 4.844963 | 0.92 | 0.04 | 0 | 12 |
| cahz | 0 | 4.839412 | 0.993 | 0.119 | 0 | 12 |
| icn | 0 | 4.645766 | 0.986 | 0.053 | 0 | 12 |
| zgc:195173 | 0 | 4.495464 | 0.957 | 0.082 | 0 | 12 |
| slc18a3a | 0 | 4.00923 | 0.829 | 0.019 | 0 | 13 |
| mdkb | 1.4E-194 | 3.845208 | 0.996 | 0.388 | 2.7E-190 | 13 |
| sox2 | 0 | 3.819129 | 0.72 | 0.029 | 0 | 13 |
| rnd3b | 0 | 3.745552 | 0.463 | 0.015 | 0 | 13 |
| kiaa0040 | 0 | 3.535853 | 0.78 | 0.038 | 0 | 13 |
| rgs3a | 2.1E-155 | 3.349586 | 0.813 | 0.219 | 4E-151 | 13 |
| arl4ab | 2.3E-244 | 3.303692 | 0.756 | 0.11 | 4.4E-240 | 13 |
| syt1a | 6E-224 | 3.09087 | 0.992 | 0.24 | 1.1E-219 | 13 |
| slit2 | 5.3E-155 | 2.999045 | 0.455 | 0.056 | 1E-150 | 13 |
| zgc:101840 | 3.6E-248 | 2.947038 | 0.679 | 0.082 | 6.7E-244 | 13 |
| icn | 0 | 4.772532 | 0.922 | 0.065 | 0 | 14 |
| sepp1a-1 | 2E-186 | 4.273613 | 0.766 | 0.079 | 3.9E-182 | 14 |
| rlbp1a | 1.7E-274 | 4.048091 | 0.906 | 0.074 | 3.2E-270 | 14 |
| sepp1a | 1.8E-170 | 4.039914 | 0.664 | 0.063 | 3.4E-166 | 14 |
| fxyd6l | 4.2E-150 | 3.916246 | 0.906 | 0.154 | 8E-146 | 14 |
| cahz | 3E-162 | 3.776796 | 0.891 | 0.131 | 5.6E-158 | 14 |
| zgc:153704 | 1.8E-205 | 3.651931 | 0.758 | 0.067 | 3.4E-201 | 14 |
| cdo1 | 7E-176 | 3.524638 | 0.742 | 0.076 | 1.3E-171 | 14 |
| LOC101885164 | 3.46E-66 | 3.308374 | 0.844 | 0.356 | 6.54E-62 | 14 |
| s100a10b | 3.6E-215 | 3.128883 | 0.742 | 0.06 | 6.9E-211 | 14 |
| fabp7a | 1.8E-162 | 4.383033 | 0.935 | 0.135 | 3.4E-158 | 15 |
| her4.3 | 9.6E-222 | 4.014462 | 0.274 | 0.006 | 1.8E-217 | 15 |
| her15.1 | 1.5E-184 | 3.997166 | 0.419 | 0.02 | 2.9E-180 | 15 |
| hmgn2 | 3.45E-46 | 3.862143 | 0.903 | 0.425 | 6.51E-42 | 15 |
| LOC100534909 | 5.2E-188 | 3.837915 | 0.419 | 0.02 | 9.9E-184 | 15 |
| LOC100148329 | 2.9E-201 | 3.669825 | 0.298 | 0.009 | 5.5E-197 | 15 |
| id1 | 1.8E-142 | 3.585239 | 0.54 | 0.046 | 3.4E-138 | 15 |
| socs3a | 2.57E-68 | 3.530901 | 0.508 | 0.08 | 4.85E-64 | 15 |
| ggctb | 7.67E-32 | 3.517551 | 0.298 | 0.053 | 1.45E-27 | 15 |
| si:dkey-238o13.4 | 7.5E-169 | 3.238404 | 0.621 | 0.051 | 1.4E-164 | 15 |
| s100a10b | 2.7E-140 | 5.259996 | 0.625 | 0.062 | 5.2E-136 | 16 |
| nrgna | 3.9E-172 | 4.525266 | 0.946 | 0.131 | 7.3E-168 | 16 |

| | | | | | | |
|---|---|---|---|---|---|---|
| syt5b | 8.9E-183 | 4.411684 | 0.982 | 0.135 | 1.7E-178 | 16 |
| plk2a | 1.3E-306 | 3.71305 | 0.438 | 0.011 | 2.5E-302 | 16 |
| rdh10a | 5.7E-158 | 3.668155 | 0.634 | 0.054 | 1.1E-153 | 16 |
| LOC110438400 | 0 | 3.492214 | 0.723 | 0.034 | 0 | 16 |
| dgkaa | 0 | 3.35081 | 0.804 | 0.018 | 0 | 16 |
| rdh10a-1 | 3.5E-132 | 3.077272 | 0.464 | 0.033 | 6.6E-128 | 16 |
| plch2a | 4.7E-169 | 3.055334 | 0.866 | 0.1 | 8.9E-165 | 16 |
| uts1 | 1.11E-94 | 3.023464 | 0.348 | 0.026 | 2.09E-90 | 16 |
| chgb | 0 | 5.855756 | 0.962 | 0.042 | 0 | 17 |
| agrn | 1.28E-49 | 4.365505 | 0.387 | 0.06 | 2.42E-45 | 17 |
| bhlhe22 | 4E-238 | 4.249286 | 0.981 | 0.091 | 7.6E-234 | 17 |
| meis2b | 2E-255 | 3.44151 | 0.943 | 0.073 | 3.8E-251 | 17 |
| slc18a3a | 0 | 3.414063 | 0.821 | 0.028 | 0 | 17 |
| syt9b | 0 | 3.390629 | 0.726 | 0.026 | 0 | 17 |
| anos1a | 1.1E-249 | 3.264021 | 0.915 | 0.068 | 2E-245 | 17 |
| phactr3b | 3.5E-127 | 3.123554 | 0.953 | 0.164 | 6.6E-123 | 17 |
| kcnc3a | 6.6E-307 | 3.04103 | 0.962 | 0.06 | 1.2E-302 | 17 |
| zgc:194629 | 0 | 2.752403 | 0.396 | 0.005 | 0 | 17 |
| cd59 | 0 | 7.808436 | 1 | 0.016 | 0 | 18 |
| plp1b | 0 | 7.33091 | 1 | 0.01 | 0 | 18 |
| cldnk | 0 | 7.209105 | 1 | 0.005 | 0 | 18 |
| cd9b | 0 | 6.571579 | 1 | 0.005 | 0 | 18 |
| si:dkey-200l5.4 | 0 | 5.958872 | 0.989 | 0.004 | 0 | 18 |
| mbpa | 0 | 5.611453 | 0.989 | 0.01 | 0 | 18 |
| tuba8l3 | 0 | 5.520353 | 0.989 | 0.005 | 0 | 18 |
| si:rp71-19m20.1 | 4.6E-253 | 5.28379 | 0.989 | 0.073 | 8.6E-249 | 18 |
| cd82a | 0 | 5.214505 | 0.989 | 0.041 | 0 | 18 |
| zwi | 0 | 5.211844 | 0.989 | 0.008 | 0 | 18 |
| apoc1 | 0 | 5.782885 | 0.884 | 0.029 | 0 | 19 |
| pfn1 | 7.2E-195 | 5.074421 | 0.884 | 0.058 | 1.4E-190 | 19 |
| cd74a | 0 | 4.617333 | 0.754 | 0.013 | 0 | 19 |
| LOC110439470 | 0 | 4.073807 | 0.609 | 0.01 | 0 | 19 |
| lgals3bpb | 0 | 3.818945 | 0.609 | 0.01 | 0 | 19 |
| vmp1 | 2.21E-18 | 3.807155 | 0.377 | 0.092 | 4.18E-14 | 19 |
| lgals2a | 8E-247 | 3.522558 | 0.652 | 0.022 | 1.5E-242 | 19 |
| fabp11a | 1.1E-102 | 3.505988 | 0.406 | 0.021 | 2.08E-98 | 19 |
| cd74b | 0 | 3.487105 | 0.565 | 0.011 | 0 | 19 |
| arpc1b | 0 | 3.378654 | 0.71 | 0.009 | 0 | 19 |
| apoc1 | 0 | 7.283479 | 0.985 | 0.029 | 0 | 20 |
| cd74a | 0 | 5.993695 | 1 | 0.012 | 0 | 20 |
| lgals3bpb | 0 | 5.980537 | 0.985 | 0.008 | 0 | 20 |

| | | | | | | |
|---|---|---|---|---|---|---|
| fabp11a | 0 | 5.878762 | 0.785 | 0.019 | 0 | 20 |
| zgc:92066 | 2.7E-54 | 5.303331 | 1 | 0.381 | 5.1E-50 | 20 |
| cmklr1 | 0 | 5.14607 | 1 | 0.005 | 0 | 20 |
| vmp1 | 4.3E-166 | 5.08591 | 1 | 0.088 | 8.1E-162 | 20 |
| si:busm1-266f07.2 | 0 | 4.799624 | 0.969 | 0.013 | 0 | 20 |
| pfn1 | 5E-237 | 4.793024 | 1 | 0.057 | 9.4E-233 | 20 |
| cd74b | 0 | 4.641042 | 0.985 | 0.009 | 0 | 20 |
| si:ch211-214p16.1 | 0 | 6.635844 | 0.875 | 0.003 | 0 | 21 |
| cxcr4b | 0 | 5.404781 | 0.786 | 0.01 | 0 | 21 |
| b2m | 1.5E-104 | 5.155761 | 1 | 0.127 | 2.8E-100 | 21 |
| ccr9a | 0 | 4.939505 | 0.786 | 0.005 | 0 | 21 |
| srgn-1 | 0 | 4.828549 | 0.929 | 0.015 | 0 | 21 |
| pfn1 | 4.8E-165 | 4.485086 | 0.911 | 0.059 | 9.1E-161 | 21 |
| LOC100151049 | 0 | 4.183341 | 0.857 | 0.011 | 0 | 21 |
| si:dkey-27i16.2 | 9.8E-295 | 4.107388 | 0.714 | 0.018 | 1.8E-290 | 21 |
| ucp2 | 7.2E-258 | 4.094345 | 0.857 | 0.031 | 1.4E-253 | 21 |
| cebpb | 1.4E-108 | 4.050617 | 0.696 | 0.05 | 2.6E-104 | 21 |
| slc6a9 | 1.65E-77 | 6.020585 | 0.947 | 0.103 | 3.12E-73 | 22 |
| tubb5 | 8E-165 | 5.812786 | 1 | 0.05 | 1.5E-160 | 22 |
| chga | 5.06E-98 | 5.139301 | 0.842 | 0.06 | 9.56E-94 | 22 |
| si:dkey-238o13.4 | 3.3E-126 | 4.751125 | 0.921 | 0.055 | 6.2E-122 | 22 |
| kidins220a | 0 | 4.70627 | 0.947 | 0.003 | 0 | 22 |
| kcnip1b-1 | 9.7E-150 | 4.493498 | 0.842 | 0.038 | 1.8E-145 | 22 |
| kcnd1 | 1.5E-124 | 4.486649 | 0.868 | 0.049 | 2.8E-120 | 22 |
| anos1a | 8.2E-98 | 4.18251 | 0.921 | 0.073 | 1.55E-93 | 22 |
| rasd1 | 1.06E-77 | 4.164931 | 0.711 | 0.052 | 1.99E-73 | 22 |
| tbx3a | 1.4E-130 | 4.120598 | 0.947 | 0.056 | 2.6E-126 | 22 |

Table S3. **The proportion of cellular changes between WT and P23H dataset**

| | **V2 run** | | | | **V3 run** | | | |
|---|---|---|---|---|---|---|---|---|
| | **WT** | | **P23H** | | **WT** | | **P23H** | |
| | # | % | # | % | # | % | # | % |
| Rods | 500 | 13.1 | 211 | 4.7 | 869 | 6.4 | 501 | 3.2 |
| Cones | 311 | 8.1 | 426 | 9.5 | 1759 | 13.0 | 1138 | 7.3 |
| New rods | 56 | 1.5 | 631 | 14.1 | 150 | 1.1 | 583 | 3.8 |
| RPC | 89 | 2.3 | 442 | 9.9 | 62 | 0.5 | 341 | 2.2 |
| RPE | 71 | 1.9 | 32 | 0.7 | 1080 | 8.0 | 156 | 1.0 |

Table S4. **Parameters of V3 datasets**

| Sample | Cells Submitted to Make Library | Total Read Number | Median Reads Per Cell | Reported Cells | Total Genes detected | Reported Median Genes | Filter Parameters | Cells Left After Filter | Median nGenes after filter |
|---|---|---|---|---|---|---|---|---|---|
| P23H | 14,000-24,000 | 608,597,236 | 35,844 | 16,979 | 22,497 | 665 | 200>x<4700 | 15,511 | 669 |
| WT | 14,000-24,000 | 967,526,548 | 58,581 | 16,516 | 21,889 | 610 | 200>x<6000 | 13,552 | 684 |
